# Brca2, Pds5 and Wapl differentially control cohesin chromosome association and function

**DOI:** 10.1101/170829

**Authors:** Ziva Misulovin, Michelle Pherson, Maria Gause, Dale Dorsett

## Abstract

The cohesin complex topologically encircles chromosomes and mediates sister chromatid cohesion to ensure accurate chromosome segregation upon cell division. Cohesin also participates in DNA repair and gene transcription. The Nipped-B – Mau2 protein complex loads cohesin onto chromosomes and the Pds5 - Wapl complex removes cohesin. Pds5 is also essential for sister chromatid cohesion, indicating that it has functions beyond cohesin removal. The Brca2 DNA repair protein interacts with Pds5, but the roles of this complex beyond DNA repair are unknown. Here we show that Brca2 opposes Pds5 function in sister chromatid cohesion by assaying precocious sister chromatid separation in metaphase spreads of cultured cells depleted for these proteins. By genome-wide chromatin immunoprecipitation we find that Pds5 facilitates SA cohesin subunit association with DNA replication origins and that Brca2 inhibits SA binding, mirroring their effects on sister chromatid cohesion. Cohesin binding is maximal at replication origins and extends outward to occupy active genes and regulatory sequences. Pds5 and Wapl, but not Brca2, limit the distance that cohesin extends from origins, thereby determining which active genes, enhancers and silencers bind cohesin. Using RNA-seq we find that Brca2, Pds5 and Wapl influence the expression of most genes sensitive to Nipped-B and cohesin, largely in the same direction. These findings demonstrate that Brca2 regulates sister chromatid cohesion and gene expression in addition to its canonical role in DNA repair and expand the known functions of accessory proteins in cohesin’s diverse functions.

**Author summary:** The cohesin protein complex has multiple functions in eukaryotic cells. It ensures that when a cell divides, the two daughter cells receive the correct number of chromosomes. It does this by holding together the sister chromatids that are formed when chromosomes are duplicated by DNA replication. Cohesin also helps repair damaged DNA, and to regulate genes important for growth and development. Even minor deficiencies in some proteins that regulate cohesin cause significant human birth defects. Here we investigated in Drosophila cells how three proteins, Pds5, Wapl and Brca2, determine where cohesin binds to chromosomes, control cohesin’s ability to hold sister chromatids together, and participate in gene expression. We find that Pds5 and Wapl work together, likely during DNA replication, to determine which genes bind cohesin by controlling how far cohesin spreads out along chromosomes. Pds5 is required for cohesin to hold sister chromatids together, and Brca2 counteracts this function. In contrast to the opposing roles in sister chromatid cohesion, Pds5 and Brca2 work together to facilitate control of gene expression by cohesin. Brca2 plays a critical role in DNA repair, and these studies expand the known roles for Brca2 by showing that it also regulates sister chromatid cohesion and gene expression. *BRCA2* mutations in humans increase susceptibility to breast and ovarian cancer, and these findings raise the possibility that changes in chromosome segregation or gene expression might contribute to the increased cancer risk associated with these mutations.

## Introduction

The cohesin complex mediates sister chromatid cohesion, which ensures accurate chromosome segregation upon cell division. Cohesin consists of the Smc1 (Flybase FBgn0040283), Smc3 (FBgn0015615), Rad21 (Vtd, FBgn0260987) and SA (FBgn0020616) proteins [1–4]. Smc1, Smc3 and Rad21 form a tripartite ring that topologically encircles chromosomes, and SA interacts with Rad21. In metazoan organisms, cohesin is loaded onto chromosomes by a complex of the Nipped-B (FBgn0026401) and Mau2 (FBgn0038300) proteins starting in early G1, and sister chromatid cohesion is established during S phase. Cohesin is removed from chromosome arms by a complex of the Pds5 (FBgn0260012) and Wapl (FBgn0004655) proteins upon entry into mitosis.

Although the Pds5-Wapl complex unloads cohesin from chromosomes, Pds5 and Wapl differ in their roles in sister chromatid cohesion. Mutations in the Drosophila *pds5* gene, similar to *Saccharomyces cerevisiae PDS5* mutations [5, 6] cause loss of sister cohesion, resulting in aneuploid cells prior to death [7]. In contrast, Drosophila *wapl* mutations cause premature loss of sister cohesion only in pericentric heterochromatic regions and impair release of sister cohesion along chromosome arms [8].

Pds5 and Wapl also control cohesin chromosome binding dynamics during interphase. In vivo FRAP (fluorescence recovery after photobleaching) experiments in Drosophila mutants show that partial reduction of Pds5 dosage increases the level of cohesin stably bound to chromosomes, consistent with its role in cohesin removal, while partial loss of Wapl unexpectedly decreases the level of stable cohesin [9]. Complete removal of Wapl, however, dramatically increases both the amount of cohesin on chromosomes and its average residence time [9]. Thus the collaborative roles of Pds5 and Wapl in cohesin removal do not fully explain their in vivo effects on cohesin dynamics and sister chromatid cohesion. Based on the in vivo FRAP data, it was hypothesized that a high Pds5 to Wapl ratio favors cohesin removal, and that Pds5 and Wapl may also have roles in cohesin regulation independently of each other [9].

More recently it was discovered that Pds5 also interacts in a 2:1 ratio with the Brca2 (FBgn0050169) DNA repair protein. This complex, which lacks Wapl, participates in meiotic recombination and mitotic DNA repair [10–12]. The Pds5-Brca2 complex occurs in multiple types of Drosophila and mammalian cells, and Drosophila Brca2 also interacts with mammalian PDS5B and PDS5A [10]. Two BRC repeats in Drosophila Brca2 interact with a region of Pds5 located adjacent to Pds5 residues that interact with Wapl, facilitating dimerization of Pds5 [11, 13].

The potential roles of the Pds5-Brca2 complex in cohesin dynamics, sister chromatid cohesin and gene transcription have not been explored. We theorized that the Pds5-Brca2 complex might help explain the differing roles of Pds5 and Wapl in cohesin dynamics and sister chromatid cohesion, and thus investigated the functions of Pds5, Brca2 and Wapl in sister chromatid cohesion, cohesin binding and localization, and gene regulation in Drosophila ML-BG3-c2 (BG3) cells. We find that Brca2 and Pds5 oppose each other in sister chromatid cohesion and in binding of the SA cohesin subunit near early DNA replication origins. Pds5 elevates the ratio of SA to cohesin at origins and Brca2 decreases SA levels. Cohesin extends outward from replication origins for several kilobases to bind active genes and enhancers, with the levels decreasing with increasing distance from the origin. Pds5 and Wapl, but not Brca2, limit the distance that cohesin spreads, thereby controlling which active genes and regulatory sequences bind cohesin. Brca2, Pds5 and Wapl have similar effects on gene expression and influence expression of the same genes as Nipped-B and cohesin. Like Nipped-B, Brca2 influences gene expression in developing wings and facilitates wing growth. These studies thus reveal previously unknown roles for Brca2 in regulation of sister chromatid cohesion and gene expression and raise the possibility that the increased cancer susceptibility caused by human *BRCA2* mutations could potentially reflect changes in cell physiology beyond DNA repair deficits.

## Results

### Brca2 opposes the role of Pds5 in sister chromatid cohesion

The discovery that Pds5 interacts with Brca2 in a complex that lacks Wapl [10–12] raised the question of whether or not Brca2 plays a role in sister chromatid cohesion. RNAi-mediated depletion of Brca2 in cultured ML-DmBG3-c2 (BG3) cells indicates that it counters the role of Pds5 in establishing or maintaining cohesion. The BG3 clonal cell line derived from larval central nervous system is diploid male, allowing accurate measurement of sister chromatid cohesion defects in metaphase chromosome spreads [14]. Figure 1A shows that as expected, depletion of Pds5 (iPds5) causes precocious sister chromatid separation (PSCS) with ~70% of chromosomes showing partial or complete separation after 5 days of treatment, and Wapl depletion (iWapl) gives less than 10% PSCS, similar to mock-treated cells.

**Fig 1.**
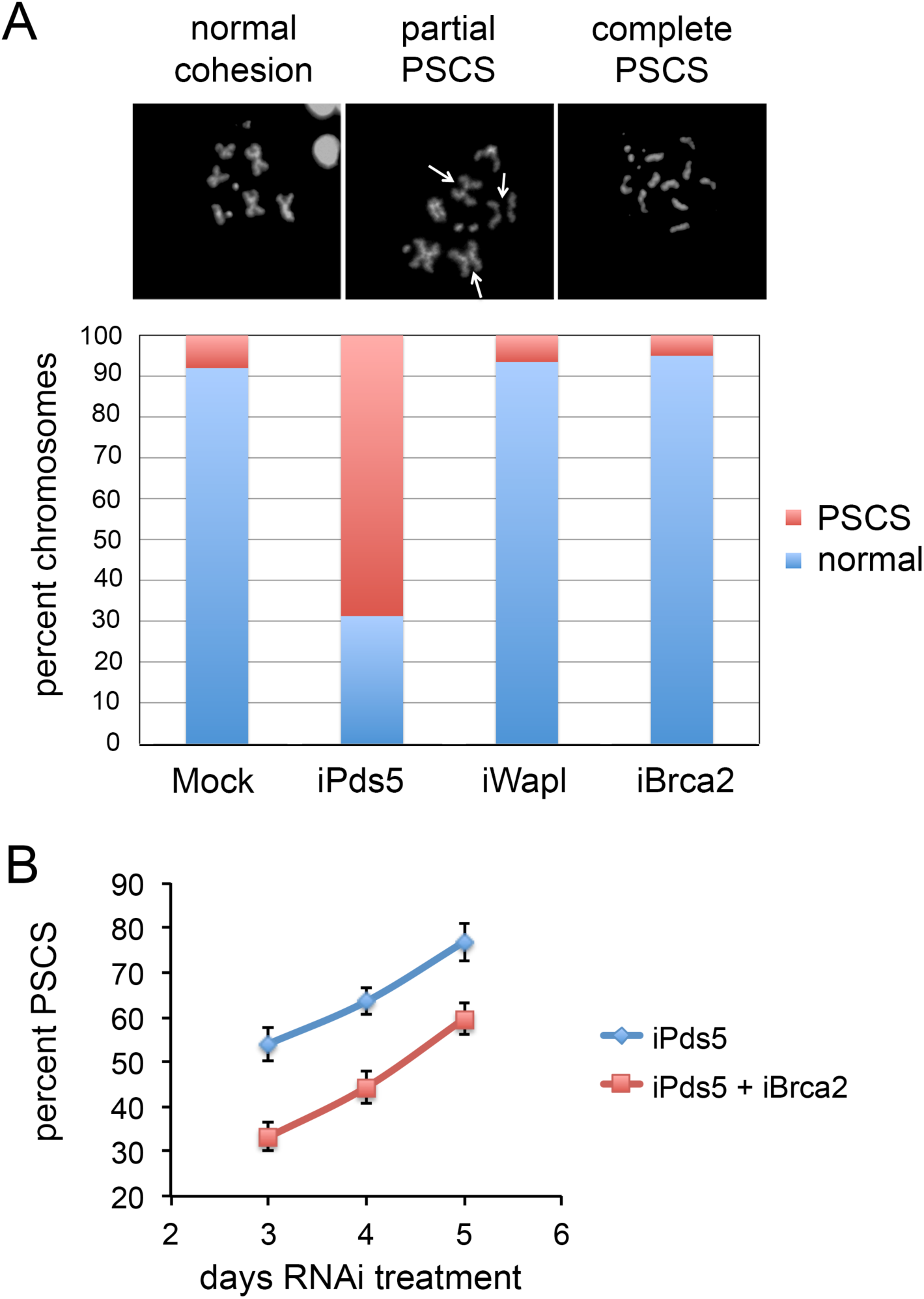
Pds5 and Brca2 oppose each other in sister chromatid cohesion in BG3 cells. (**A**) The three micrographs show examples of normal metaphase chromosomes and of precocious sister chromatid separation (PSCS). The bar graph shows the percent of chromosomes showing normal cohesion (blue) or partial or complete PSCS (red) in mock-treated (Mock), and Pds5-depleted (iPds5), Wapl-depleted (iWapl) or Brca2-depleted (iBrca2) BG3 cells after 5 days of RNAi treatment. (**B**) The graphs show the mean percentage of chromosomes showing PSCS in individual cells after three to five days of RNAi treatment for Pds5 only (iPds5, blue) or for both Pds5 and Brca2 (iPds5 + iBrca2, red). Error bars are standard errors of the mean. A minimum of 50 metaphase nuclei were scored for each individual group and time point. Similar results were obtained in two additional experiments.

Brca2 depletion (iBrca2) alone does not alter PSCS frequency (Fig 1A) but if Brca2 is co-depleted with Pds5 (iPds5 iBrca2) PSCS frequency is significantly reduced compared to Pds5 depletion alone (Fig 1B). Pds5 depletion reduces cell proliferation after 5 days of RNAi treatment, presumably reflecting increasing aneuploidy as observed in *pds5* mutant larval neuroblasts [7] while there is no discernable effect of Wapl or Brca2 depletion on cell growth. These results indicate that Brca2 opposes the role of Pds5 in sister chromatid cohesion.

Brca2 is unlikely to oppose Pds5 function indirectly through effects on the levels of other sister chromatid cohesion proteins. Western blots of whole cell extracts illustrating Pds5, Wapl and Brca2 depletion are shown in S1 Figure. These westerns also show that Pds5 depletion does not significantly alter the total levels of Nipped-B, Wapl or the cohesin subunits examined. Brca2 depletion does not increase or decrease the levels of Pds5, although Pds5 depletion modestly reduces Brca2 levels.

In RNA-seq experiments described in more detail in a later section, Pds5 depletion increases *Brca2* gene transcripts 1.4-fold (S1 Figure, panel I). Thus the partial decrease in Brca2 protein caused by Pds5 depletion likely represents destabilization of Brca2 protein, suggesting that Brca2 is stabilized by interaction with Pds5.

The RNA-seq experiments also show that Pds5 and Brca2 depletion have similar modest effects on RNA levels for some genes encoding sister chromatid cohesion proteins, and indicate that Pds5-Brca2 co-depletion does not suppress sister chromatid cohesion defects by altering expression of these genes. Prior studies show that 80 to 90% depletion of Nipped-B or the SA or Rad21 cohesin subunits in BG3 cells does not measurably alter sister chromatid cohesion [14]. As shown in S1I Figure the effects on cohesion factor transcript levels are all less than 2-fold, making it unlikely that they are involved in the substantial changes in sister chromatid cohesion observed in the Pds5 depletion and their suppression in the Pds5-Brca2 co-depletion. Pds5 and Brca2 depletion both modestly reduce *Nipped-B* RNA levels by 30 to 40%, and increase transcripts of the *Smc1*, *Smc3*, *wapl*, *eco*, and *dmt* (*dalmatian*, encoding a sororin-shugoshin fusion protein [15, 16]) gene up to 1.5-fold, none of which explains the opposing effects of Pds5 and Brca2 on sister cohesion. The only significant effect of the double Pds5-Brca2 depletion that is not observed in the single depletions is a 40% reduction in *SA* transcripts, which should reduce, not rescue cohesion if it were of sufficient magnitude to actually have an effect (S1I Fig). No other known cohesion factor gene transcript levels are significantly affected by Pds5 and/or Brca2 depletion (S1I Fig).

Taken together, the lack of substantial effects of Pds5 and/or Brca2 depletion on the levels of sister chromatid cohesion proteins assayed and all known cohesion factor transcripts indicate that the suppression of the cohesion defects in Pds5-Brca2 co-depleted cells relative to cells depleted for Pds5 is unlikely to be caused by changes in the levels of sister chromatid cohesion proteins. This supports the idea that Brca2 directly opposes the role of Pds5 in sister chromatid cohesion through its interaction with Pds5. We cannot, however, eliminate the possibility of indirect effects through unknown cohesion proteins.

### Pds5 and Brca2 chromosome binding patterns differ from cohesin and Wapl

To gain insights into how Pds5, Wapl and Brca2 influence cohesin function and sister chromatid cohesion we mapped them genome-wide in BG3 cells by chromatin immunoprecipitation with high-throughput sequencing (ChIP-seq). As described elsewhere for Nipped-B and cohesin [17, 18] we used at least three biological replicates for each protein, sequencing each replicate to a minimum of 10X genome coverage, normalizing each to input chromatin sequenced to at least 45X genome coverage. In Drosophila, cohesin and Nipped-B occupancy spread broadly over several kilobases, and ultra-deep sequencing and calculating enrichment relative to input chromatin instead of calling peaks are vital for accurate and reproducible results [18]. Enrichment relative to input chromatin was calculated using 250 bp sliding windows to generate values every 50 base pairs, thereby reducing noise and facilitating direct comparisons of replicates and downstream calculations. The biological replicates showed high genome-wide Pearson correlations (0.65 to 0.9) and were averaged for further analysis. Preimmune serum precipitations show no regions with significant enrichment using these procedures, indicating an absence of methodological artifacts. S2A Figure shows examples of the correlations between three SA ChIP-seq biological replicates and S2B Figure shows a genome browser example of ChIP-seq with preimmune serum compared to multiple ChIP-seq experiments.

We compared the Pds5, Wapl and Brca2 binding patterns to the Rad21 cohesin subunit, and prior ChIP-seq data [17] for the SA cohesin subunit and Nipped-B. As illustrated by the *kayak* gene region in Figure 2A, Wapl displays a broad spreading pattern nearly indistinguishable from cohesin. Nipped-B differs somewhat from cohesin and Wapl with noticeable broad peaks superimposed on the spreading pattern. Pds5 and Brca2 show substantially less spreading, being more restricted to broad peaks that co-localize with Nipped-B peaks (Fig 2A).

**Fig 2.**
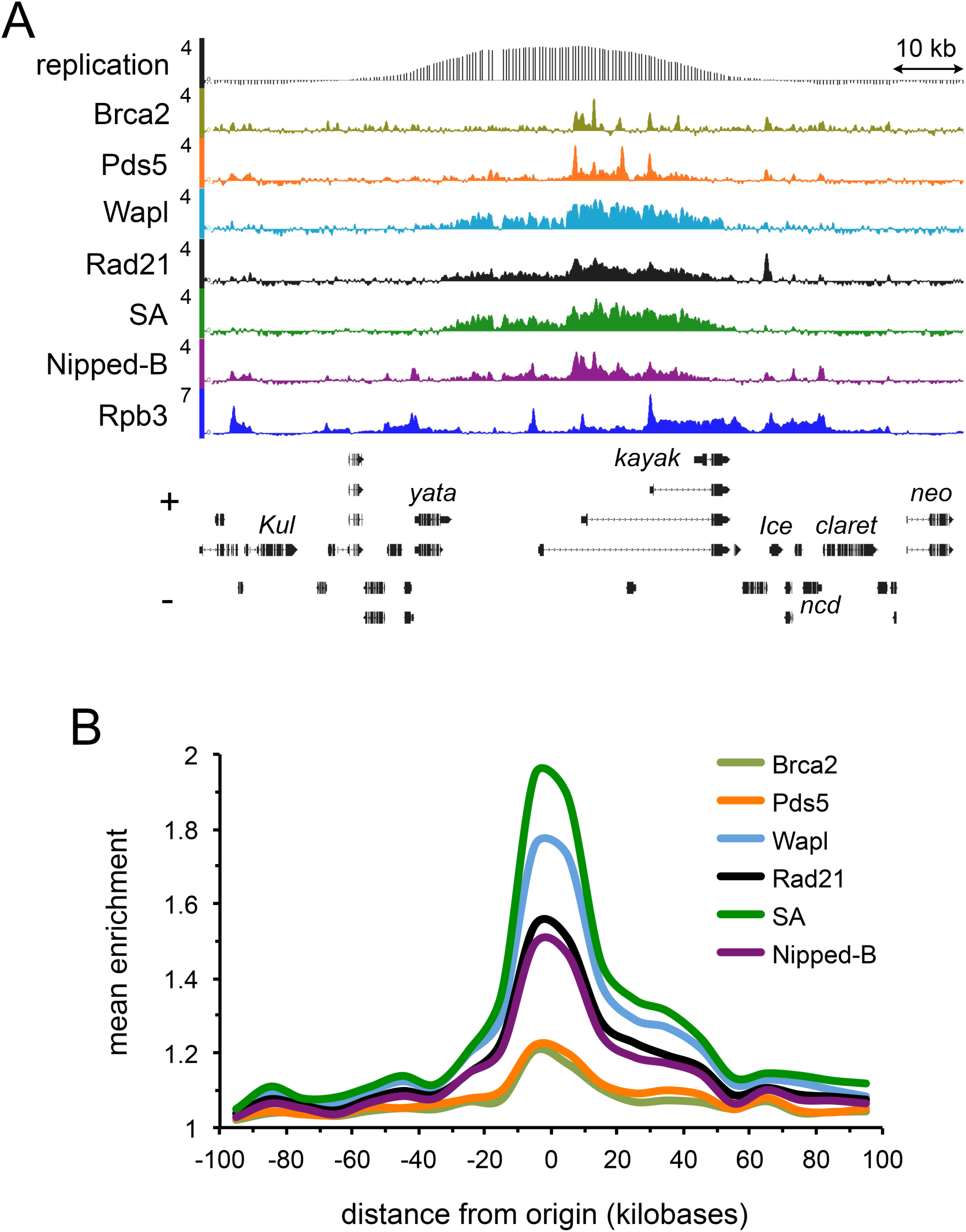
Cohesin and the accessory factors localize to DNA replication origins in BG3 cells. (**A**) Genome browser view of Brca2, Pds5, Wapl, Rad21, SA, Nipped-B and Rpb3 ChIP-seq at the *kayak* locus containing an early DNA replication origin. The scales are log2 enrichment. David MacAlpine provided the processed BG3 early DNA replication data (GEO accession GSE17287) and the scale is MA2C score. The SA, Nipped-B, and Rpb3 ChIP-seq data are published elsewhere [17, 33]. (**B**) Meta-origin analysis of the ChIP-seq data using the 78 strongest early DNA replication origins (positions in S3 File). The mean enrichment in 10 kb bins was calculated from −100 kb to +100 kb from the origin centers.

The restricted Pds5 localization relative to Wapl was unexpected given that Pds5 and Wapl form a complex and can bind together to cohesin [19–21]. Our findings indicate that Wapl also binds cohesin independently of Pds5 in a location-dependent manner. Independent Wapl binding is consistent with the finding the N terminal region of human Wapl binds to the C terminal region of Rad21 in the presence of SA and absence of Pds5 [22]. The differences in Pds5 and Wapl localization also predict that the dynamics of cohesin removal differs along the chromosome. Interaction of Wapl with the N terminal region of Rad21 requires Pds5 [22]. This interaction is required for the Pds5-Wapl complex to open the Rad21 N terminus interface with the Smc3 ATPase head domain and remove cohesin from chromosomes [2, 13, 23–25]. Thus cohesin is less likely to be removed in the regions with high Wapl and low Pds5. This idea is also consistent with in vivo FRAP data with Drosophila *pds5* and *wapl* mutants showing that a low Pds5 to Wapl ratio increases cohesin binding, and that a high ratio decreases binding [9].

Pds5 and Brca2 occupancy overlap with each other, as would be expected if they interact with each other on chromosomes. Using a threshold of enrichment in the 95^th^ percentile or greater over regions ≥300 bp in length to call binding, there are 6,452 Pds5 euchromatic sites and 6,430 Brca2 euchromatic sites of variable length, 3,600 (56%) of which overlap. The actual overlap is greater because many sites show significant enrichment for the other protein below the 95^th^ percentile. Examples of binding regions called at the 95^th^ percentile are shown in S2B Figure. Visual inspection in a browser reveals a minor number of sites enriched only for Brca2 or for Pds5, which are typically short regions with low enrichment (S2B Fig, asterisks). The vast majority of Brca2 and Pds5 binding sites occur within broad regions occupied by cohesin and Wapl, with a few exceptions of small peaks with lower enrichment (S2B Fig, daggers).

### Sister chromatid cohesion factor occupancy centers at early DNA replication origins

Visual inspection in a browser revealed that most sites with high Pds5 and Brca2 occupancy in BG3 cells are located within a few kilobases of early DNA replication origins. Sister chromatid cohesion is established during DNA replication, and cohesin is enriched and loaded at replication origins and sites in Drosophila, yeast and vertebrates [26–32]. We thus conducted an origin meta-analysis of all cohesion factor ChIP-seq data, finding that the highest Nipped-B, cohesin, Pds5, Wapl and Brca2 levels all center at origins. The locations of early DNA replication origins in BG3 cells were reported by the modENCODE project and confirmed by mapping pre-replication complex (pre-RC) binding sites [26]. As shown in Figure 2A, a replication origin is located within multiple active genes that bind RNA polymerase II (Rpb3) [33] in the *kayak* region. We averaged the ChIP-seq enrichment for cohesin and accessory factors in non-overlapping 10 kb bins from −100 kb to +100 kb outward from the centers of the 78 strongest early origins in BG3 cells (positions in S3 File). As revealed in Figure 2B, all show maximal occupancy near the metaorigin center, with the levels decreasing outward in both directions.

The ChIP-seq experiments were conducted with asynchronously dividing cells, and although the cohesion factor binding centers at replication origins, most cells (~85%) are in interphase, in which cohesin binds specifically to active genes and enhancers [34, 35]. Thus the metaorigin analysis reveals that the further an active gene is from the origin, the less likely it is to bind cohesin. For instance, *kayak* under the origin center binds cohesin, and *yata* some 20 kb from the origin center binds little, even though it is transcribed (Fig 2A). Inactive genes and intergenic regions located between an active gene that binds cohesin and the origin do not bind cohesin. Thus cohesin binding is discontinuous under most origins and in the origin-flanking regions, with active genes close to the origin binding high cohesin levels, and genes further away binding less or none. The cohesin pattern revealed by the metaorigin analysis thus does not directly measure binding to origins per se, but shows that origins have a strong positive influence on cohesin binding to active genes in their vicinity.

Prior studies show that genes whose expression and transcription are altered by Nipped-B or cohesin are enriched for those that bind high levels of cohesin [34, 35]. Because the metaorigin analysis indicates that genes located closer to origins bind higher levels of cohesin than those further away, we can deduce that genes positioned closer to origins are more likely to be altered in transcription with changes in cohesin dosage than are genes positioned further away. It is important to remember, however, that many genes that bind high levels of cohesin show little change in expression upon cohesin depletion, and that many others that bind little or no cohesin are affected indirectly [34, 35]. Thus for any individual gene, their distance from an origin cannot predict the effect of cohesin on their transcription.

### Pds5 and Wapl control cohesin localization around replication origins

Pds5, Brca2 and Wapl all have different effects on sister chromatid cohesion. Sister chromatid cohesion is established during DNA replication, and above we show that binding of these regulatory factors centers at early replication origins. This led us to hypothesize that Pds5, Wapl and Brca2 differentially affect sister chromatid cohesion because they differentially influence cohesin binding at origins. We thus compared their effects on cohesin binding by metaorigin analysis. Cohesin ChIP-seq was performed for BG3 cells depleted for Pds5, Brca2 or Wapl using multiple biological replicates for each depletion. As noted above, depletion of each of these proteins has no significant effects on the levels of each other than a modest decrease in Brca2 with Pds5 depletion (S1 Fig). Also as noted above, Wapl and Brca2 depletion have no measurable effect on cell division, but Pds5 depletion, which causes loss of sister chromatid cohesion, causes cells to stop dividing after five days of RNAi treatment, so chromatin was isolated prior to five days.

These experiments revealed that depletion of Pds5 or Wapl, but not Brca2, increases the distance that the domains of cohesin occupancy extend outward from origins. Figure 3A shows that Pds5 depletion (iPds5) increases spreading of the Rad21 and SA cohesin subunits, Wapl, and Nipped-B in the regions flanking the *kayak* replication origin. Metaorigin analysis of the Rad21 data shows that this effect is global, with statistically significant increases in occupancy for tens of kilobases in the flanking regions (Fig 3B). Wapl depletion (iWapl) similarly increases cohesin spreading around origins as illustrated for the SA cohesin subunit at *kayak* locus in Figure 4A and metaorigin analysis (S4A Fig). Brca2 depletion (iBrca2) does not increase the size of cohesin domains (Fig 4A). Although Pds5 depletion reduces chromatid cohesion, Wapl depletion does not. We thus conclude that the expansion of cohesin territories does not reflect loss of sister cohesion. Because Wapl depletion has no measurable effect on cell division, which is reduced by Pds5 depletion, we also conclude that changes in the cell cycle do not cause the cohesin domain expansion.

**Fig 3.**
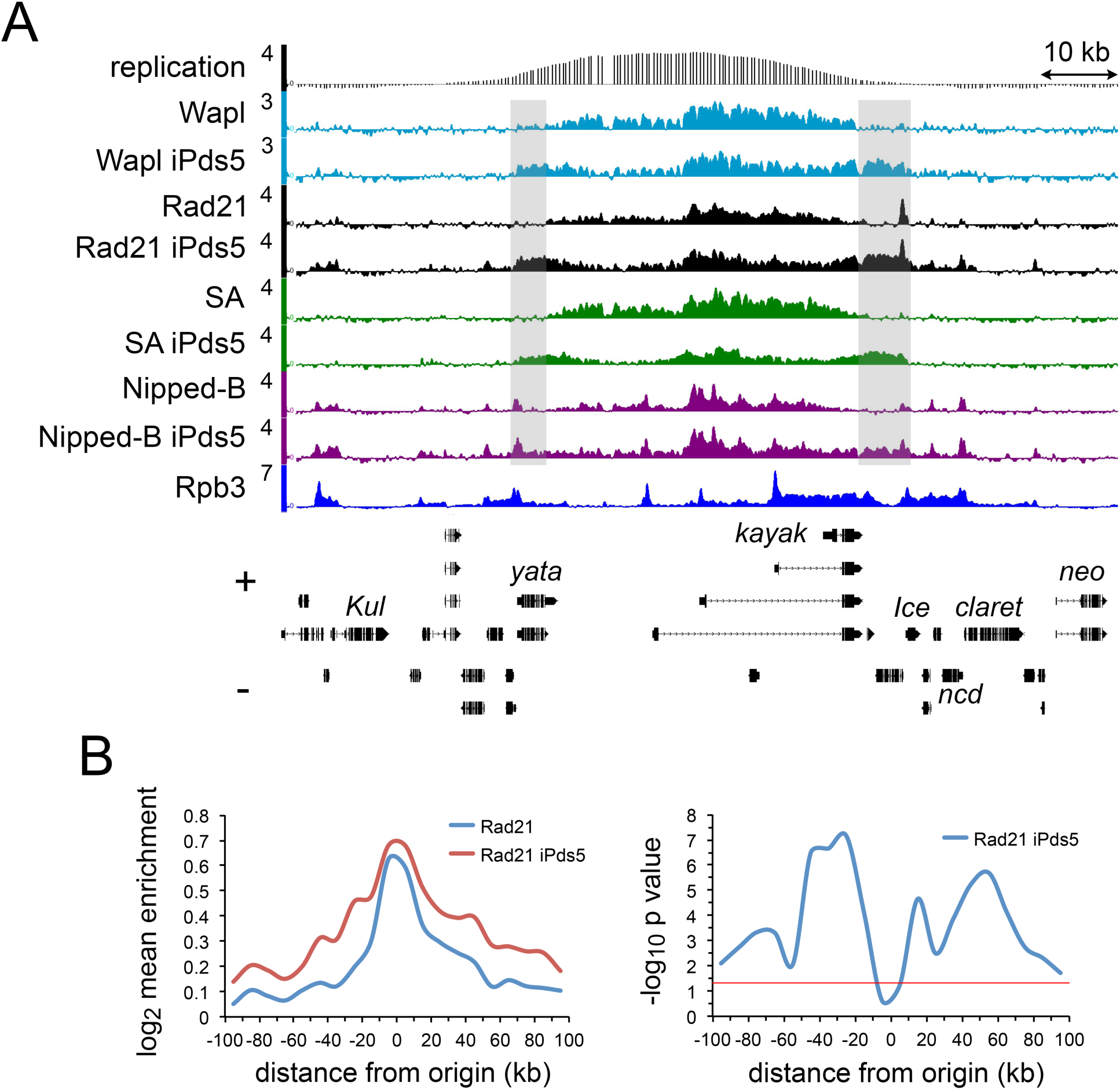
Depletion of Pds5 causes extension of cohesin and accessory factor binding domains surrounding replication origins. (**A**) Genome browser view of the *kayak* origin region, showing ChIP-seq for Wapl, Rad21, SA, and Nipped-B in control cells and cells depleted for Pds5 (iPds5). Shaded areas show regions with increased binding of cohesin, Wapl and Nipped-B in Pds5-depleted cells. (**B**) The left panel shows the Rad21 meta-origin analysis in mock control cells (Rad21, blue) and cells depleted for Pds5 (Rad21 iPds5, red). The right panel shows the –log10 p values for differences in Rad21 enrichment in each bin used for the meta-origin analysis. P values were calculated using the Wilcoxon signed rank test.

**Fig 4.**
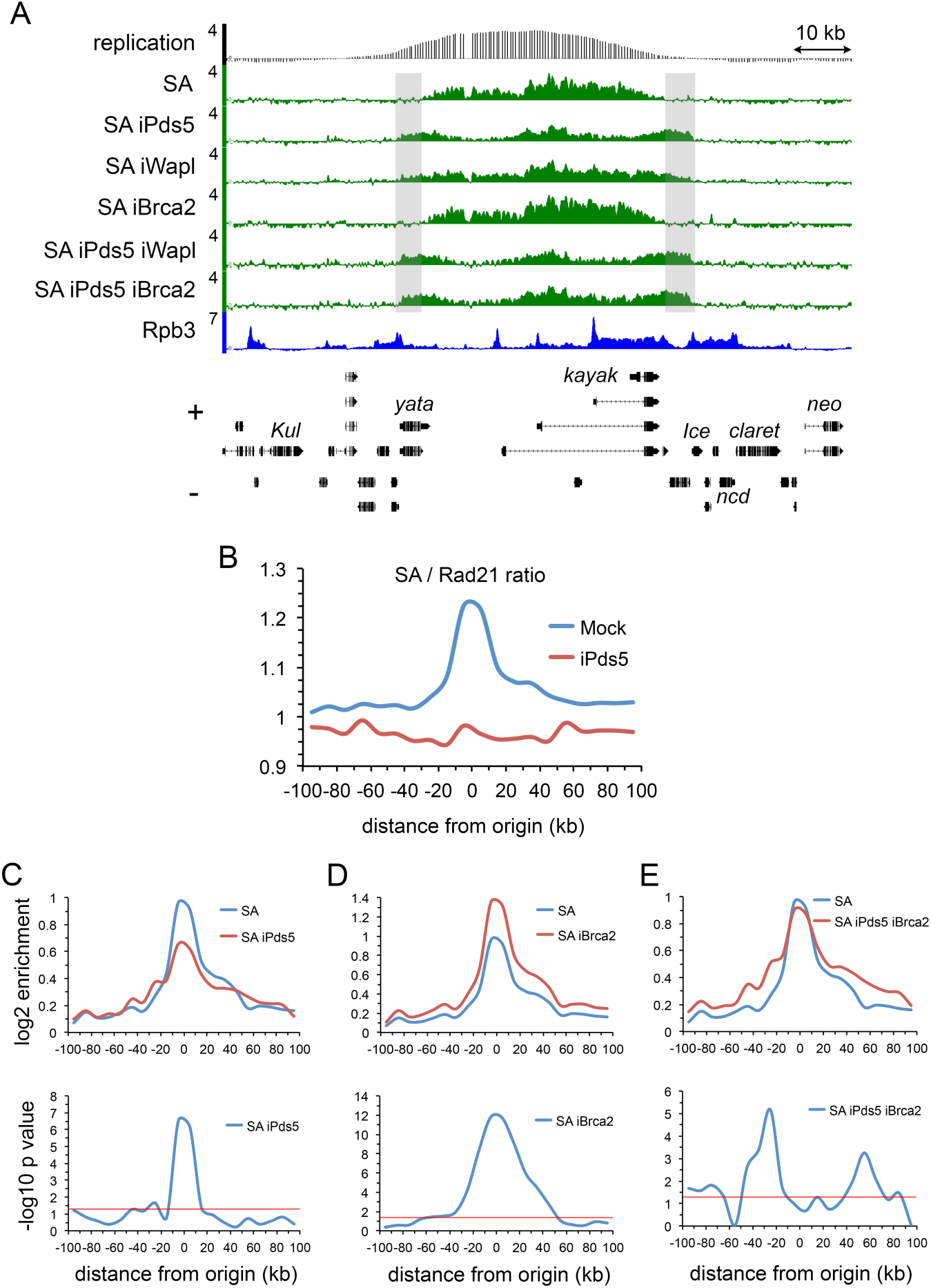
Pds5 and Brca2 have opposing effects on SA cohesin subunit levels at DNA replication origins. (**A**) Genome browser view of SA ChIP-seq at the *kayak* locus in mock-treated BG3 cells (SA) and BG3 cells RNAi depleted for Pds5 (iPds5), Wapl (iWapl), Brca2 (iBrca2), Pds5 and Wapl (iPds5 iBrca2) and Pds5 and Brca2 (iPds5 iBrca2). (**B**) Metaorigin analysis of the SA to Rad21 ChIP-seq enrichment ratio in mock control cells (Mock, blue) and cells depleted for Pds5 (iPds5, red). (**C**) The top panel is the metaorigin plot of SA enrichment in control (SA, blue) and Pds5-depleted cells (SA iPds5, red). The bottom panel is the meta-origin plot of –log10 p values for the difference in SA enrichment calculated using the Wilcoxon signed rank test. (**D**) Same as C except for cells depleted for Brca2 (iBrca2). An example of the increase in SA at the kayak locus is shown in S2C Figure. (**E**) Same as C with cells depleted for both Pds5 and Brca2 (iPds5 iBrca2).

The expansion of cohesin territories with Pds5 depletion indicates an increase in the total amount of cohesin bound to chromosomes, consistent with in vivo cohesin FRAP data in heterozygous *pds5* mutants [9]. Metaorigin analysis, however, indicates that the increase is regional, occurring primarily at locations more distant from replication origins. The increase in cohesin at these locations reflects association of cohesin with active genes that normally bind little or no cohesin, such as *yata* in Figure 4A. Active genes bind cohesin, and inactive genes or intergenic regions do not, and this holds true for the extended cohesin domains with Pds5 depletion. Skipping of the extended cohesin domains over inactive genes and intergenic regions is illustrated at the *string* locus in S5A Figure. These findings indicate that many active genes have the potential to bind cohesin, but normally do not because they are too far from an origin.

Cohesin domain expansion upon Pds5 or Wapl depletion may be linked to changes in chromosome architecture. The ends of the cohesin domain at the Enhancer of split gene complex coincide with the borders of a topologically-associating domain (TAD) [36]. Pds5 or Wapl depletion causes cohesin to extend beyond these borders (S5B Fig). Thus Pds5 or Wapl depletion might alter the locations of the TAD boundaries, or alternatively, reduce the abilities of the boundaries to block cohesin spreading. DNA loop extrusion through cohesin-CTCF insulator complexes can form TADs in mammalian cells and Wapl restricts loop extrusion [37] but a similar mechanism appears unlikely at Enhancer of split where TAD structure is independent of cohesin or insulator proteins, as shown by cohesin depletion and the lack of CTCF or other insulator proteins near the boundaries [36]. Many other Drosophila TADs also form independently of cohesin or CTCF [38–40].

The finding that active genes closer to replication origins are more likely to bind cohesin lead us to theorize that DNA replication helps determine which genes bind cohesin. DNA replisomes push topologically-bound cohesin in single molecule studies in Xenopus extracts [41] suggesting that cohesin loaded at origins may be pushed outward by replication forks, and data in yeast suggest that cohesin loaded at replication sites travels with replication forks [32]. We reasoned that if DNA replication pushes cohesin, then the cohesin domain expansion caused by Pds5 or Wapl depletion could reflect either a reduced rate of cohesin removal in front of replication forks, or increased speed of replication fork movement. DNA fiber assays, however, show that Pds5 depletion does not alter fork speed (S6 Fig). Brca2 depletion also does not alter replication speed, but as reported for mammalian cells [42] increases degradation of newly-replicated DNA behind stalled forks (S6 Fig). Thus increased cohesin spreading does not reflect an increase in replication fork speed, leading us to theorize that it may be caused by a reduced rate of cohesin removal in front of replication forks.

### Pds5 and Brca2 oppositely regulate SA cohesin subunit levels at DNA replication origins

Metaorigin analysis unexpectedly revealed that the SA to Rad21 cohesin subunit ratio is elevated at origin centers compared to flanking regions. The ratio of the mean SA enrichment to mean Rad21 enrichment was calculated for each 10 kb bin for each early origin, and the average ratio for each bin across all origins is plotted in Figure 4B. This is equivalent to normalizing enrichment for a histone modification to total histone occupancy, or a Pol II modification to total Pol II [33]. The high SA to Rad21 enrichment ratio indicates that the SA to Rad21 stoichiometry is higher near origins. The enrichment ratio does not measure actual stoichiometry, or the exact extent to which stoichiometry differs, but indicates the direction in which it varies or changes. It is possible that some or all of the increased SA at origins binds independently of the cohesin complex.

Strikingly, Pds5 depletion reduces SA levels at origin centers (Fig 4C) and equalizes the SA to Rad21 ratio across the metaorigin and flanking regions (Fig 4B). Equalization of the SA to Rad21 ratio across the entire 200 kb metaorigin region upon Pds5 depletion indicates that at origins, Pds5 actively increases SA levels above a default SA to cohesin ratio. In contrast, Wapl depletion does not alter SA levels at origins (S4A Fig) and Brca2 depletion substantially increases SA occupancy in the metaorigin analysis (Fig 4D) and as illustrated specifically at the *kayak* locus (S2C Fig). When Pds5 and Brca2 are co-depleted, their opposite effects cancel out, leaving little net change in SA levels at origin centers (Fig 4E). This makes the expansion of the SA domains from origins more evident (Fig 4E) giving a pattern similar to that observed with Rad21 (Fig 3B) with statistically significant increases over several kilobases in the flanking regions. Combined, these findings indicate that Pds5 actively increases SA levels at origins, and that Brca2 counters this Pds5 activity.

The opposing roles of Pds5 and Brca2 in controlling SA occupancy at origins parallels their opposing effects on sister chromatid cohesion. The tripartite Smc1-Smc3-Rad21 ring topologically binds chromosomes in the absence of SA or Pds5 [43] both of which are required for sister chromatid cohesion. We thus speculate that Pds5 and Brca2 regulate the SA to cohesin ratio at origins to determine the fraction of cohesin complexes that becomes cohesive behind replication forks. This is consistent with the idea that SA can link two cohesin rings to establish cohesion via a handcuff mechanism [44, 45]. An alternative idea is that SA might facilitate cohesin topological binding around both newly-synthesized sister chromatids behind replication forks [43]. We cannot measure the actual SA to Rad21 stoichiometry at various points along the chromosome, and thus do not know if some cohesin rings lack SA, as some of these ideas suggest. However, we can conclude that the role of Pds5 in increasing SA at replication origins differs from its global effect of reducing total cohesin binding revealed by in vivo FRAP experiments [9] and the Rad21 metaorigin analysis. We thus also theorize that proteins that bind specifically to replication origins modify Pds5 activity, or that this particular Pds5 activity is limited to early G1 when origins are licensed, or early S phase when origins fire.

### Pds5 depletion increases Nipped-B levels at origins and extends Nipped-B and Wapl binding domains

Other findings support the idea that Pds5 activity at origins differs from other locations. We considered the possibility that some of the differential effects of Pds5, Wapl, and Brca2 depletion on cohesin levels and distribution around origins could reflect how these factors influence each other’s association with chromosomes, or binding of the Nipped-B cohesin loader. As shown in S1 Figure, other than a modest decrease in Brca2 with Pds5 depletion, depletion of these factors do not significantly alter the total levels of each other. As detailed below, we find that the cohesin accessory factors influence each other’s association with chromosomes, but the changes do not result in the expected effects on cohesin levels at origins, supporting the idea that their activities at origins differ from their roles in the flanking regions.

The effect of Pds5 depletion on Wapl chromosome binding parallels the effects it has on SA. The Wapl to Rad21 ratio, like the SA to Rad21 ratio, is higher at origin centers than in flanking regions, and Pds5 depletion equalizes this ratio across the origin and flanking regions (Fig 5A). Pds5 depletion does not noticeably alter total Wapl proteins levels (S1 Fig). Similar to the effect on SA, Pds5 depletion decreases Wapl at origins with moderate increases in flanking regions (Fig 5B). These results indicate that although Pds5 facilitates Wapl binding at origins, it is not essential for Wapl binding. In contrast, Pds5 depletion has only minor effects on Brca2 levels at origins and flanking regions (Fig 5C) although total Brca2 protein levels are modestly reduced (S1 Fig). Thus Pds5 is not required for Brca2 to bind to chromosomes.

**Fig 5.**
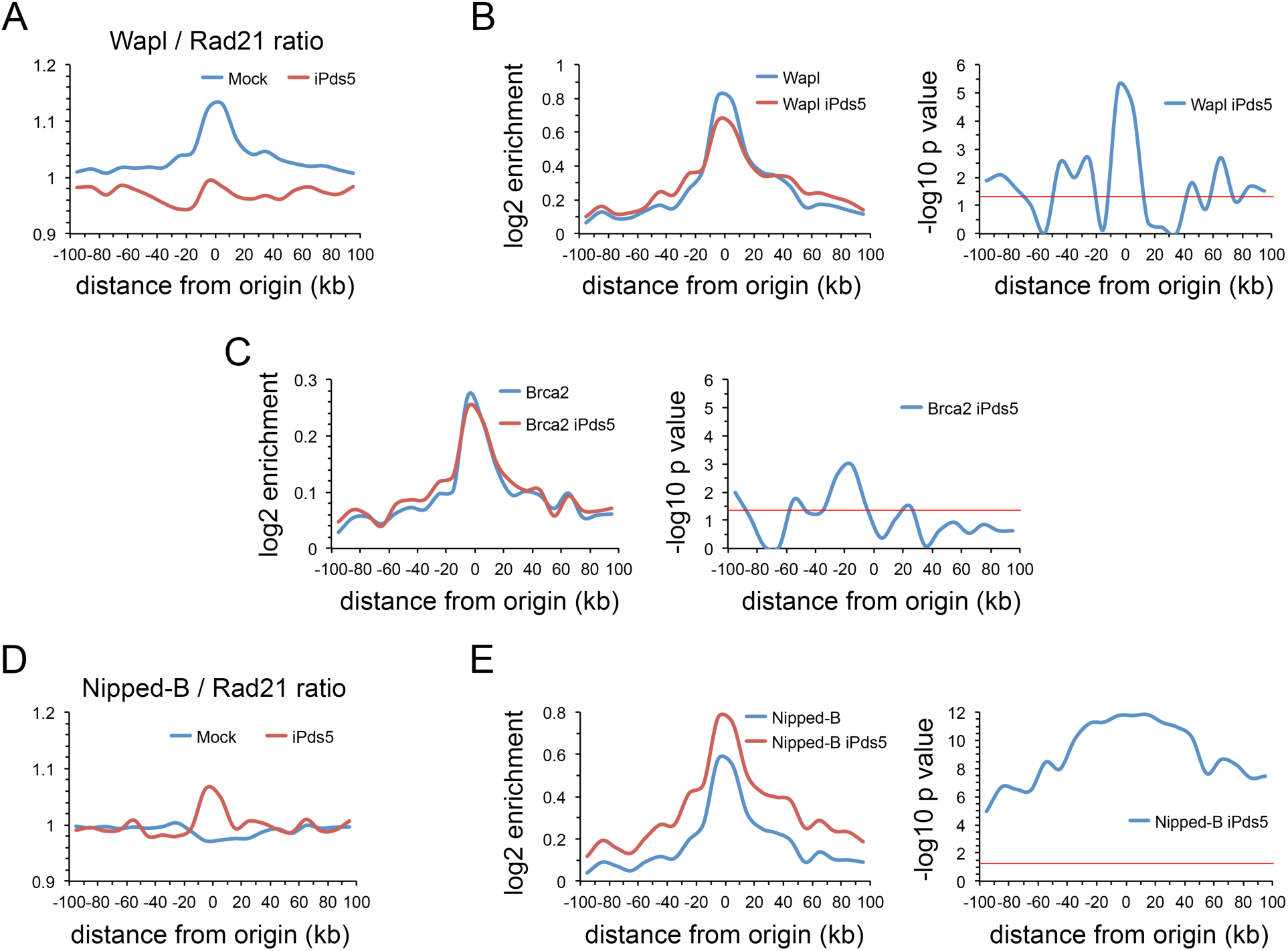
Pds5 influences Wapl and Nipped-B binding at DNA replication origins with little effect on Brca2. (**A**) Metaorigin plot of Wapl to Rad21 ratio in control cells (Mock, blue) and cells depleted for Pds5 (iPds5, red). (**B**) Left panel is meta-origin analysis of Wapl ChIP-seq enrichment in control cells (Wapl, blue) and cells depleted for Pds5 (Wapl iPds5, red). Right panel is the plot of – log10 p values for the differences in Wapl enrichment in the meta-origin bins calculated using the Wilcoxon signed rank test. (**C**) Same as B for Brca2 enrichment. (**D**) Same as A for the Nipped-B to Rad21 ratio. (**E**) Same as B for Nipped-B ChIP-seq enrichment.

Because the Pds5-Wapl complex removes cohesin from chromosomes, depletion of Pds5 with the accompanying decrease in Wapl should increase cohesin levels at origins, but as shown above, SA decreases (Fig 4C) and Rad21 is not altered (Fig 3B). Thus Pds5 and Wapl together do not effectively remove cohesin at origins, supporting the idea that their activities differ at origins than in other regions.

The Nipped-B to Rad21 ratio is constant across the 200 kb metaorigin region, but increases at the origin upon Pds5 depletion, accompanied by an overall increase in Nipped-B levels across the metaorigin (Fig 5D,E). Pds5 depletion does not alter total Nipped-B protein levels (S1 Fig). We hypothesize that Pds5 and Nipped-B, which have very similar 3D HEAT repeat structures [13, 46–50] compete for binding to cohesin or other proteins at origins, and that loss of Pds5 allows more Nipped-B to bind. This idea is supported by in vivo FRAP studies in which Nipped-B overexpression in the absence of excess Mau2 increases global cohesin binding and nearly doubles cohesin’s chromosomal residence time, consistent with a decrease in Pds5 activity [9]. Biochemical experiments show that the Nipped-B and Pds5 orthologs in *C thermophilium* bind to overlapping regions in Rad21, further supporting this idea [47].

Higher Nipped-B occupancy caused by Pds5 depletion, similar to loss of Pds5, should also increase cohesin levels at origins, but cohesin increases occur only in the flanking regions, not at origin centers. This lends further support to the idea that cohesin accessory factor activity is modified at origins.

Wapl depletion increases Pds5 (S4B Fig) and decreases Nipped-B (S4C Fig) at origins although the total levels of Pds5 and Nipped-B proteins are not altered (S1 Fig). Based on the FRAP [9] and biochemical [47] evidence that Pds5 and Nipped-B compete for cohesin, we posit that Wapl may facilitate Nipped-B binding and that loss of Nipped-B upon Wapl depletion permits increased Pds5 occupancy. Alternatively, Wapl could inhibit Pds5 binding directly and the increase in Pds5 caused by Wapl depletion could indirectly decrease Nipped-B binding.

Importantly, the increase in Pds5 upon Wapl depletion shows that Wapl does not recruit Pds5 to chromosomes. Instead, Brca2 is more crucial, as Brca2 depletion significantly decreases Pds5 occupancy (S4D Fig) although there is no significant effect on total Pds5 protein levels (S1 Fig). In contrast, Brca2 depletion has little effect on Wapl occupancy (S4E Fig). Nipped-B and Rad21 depletion also decrease Pds5 binding (S4F,G Fig) indicating that interaction of Pds5 with cohesin is also important for Pds5 binding to chromosomes, consistent with prior studies [5, 6, 51, 52].

The key finding that emerges from the above experiments is that, in contrast to the effects on cohesin occupancy in origin-flanking regions, the effects of Pds5 and Wapl depletion on cohesin levels at origins are not explained by their roles in removing cohesin from chromosomes, or their effects on the Nipped-B loading factor. We thus hypothesize that other factors, potentially DNA replication proteins such as the origin recognition complex (ORC) or the MCM helicase complex, modify the activities of cohesin regulatory factors at origins. It is also possible that their activities are altered in early G1, when origins are licensed, or early S phase, when the origins fire.

### The ratios of accessory factors to cohesin vary between different types of gene regulatory sequences

The above analysis shows that Pds5 and Wapl globally determine which active genes and regulatory sequences bind cohesin by limiting how far cohesin domains extend outward from replication origins, and that Pds5 and Brca2 control the ratio of SA to cohesin at origins. The metaorigin big picture view, however, does not indicate if association of cohesin and its regulatory factors differ between different types of gene regulatory sequences in the cohesin-binding domains, or how depletion of these factors might influence cohesin binding to different types of gene regulatory elements. Nipped-B and cohesin preferentially bind active promoters with high levels of promoter-proximal pausing, and essentially all transcriptional enhancers and Polycomb Response Elements (PREs) [14, 17, 33–36, 53, 54]. Although the metaorigin analysis indicates that proximity to a replication origin determines the likelihood that an active gene or regulatory sequence will bind cohesin, variations in the levels of cohesin and it’s accessory factors at different types of regulatory sequences could potentially impact interphase cohesin dynamics and gene expression.

We find that the relative levels of the different cohesin regulatory factors vary substantially between active promoters, enhancers and PREs, indicating that interphase cohesin binding dynamics might differ between these types of regulatory sequences. The mean ChIP-seq enrichment for these proteins at each of the active promoters, extragenic enhancers and PREs in BG3 cells were calculated as previously described [17] and the distributions of the mean enrichments across all active promoters, enhancers and PREs are presented as violin plots in Figure 6. Active promoters, enhancers and PREs were defined as described in Figure 6, and enhancers located within transcribed regions were excluded to minimize the influence of transcription through a sequence on protein binding.

**Fig 6.**
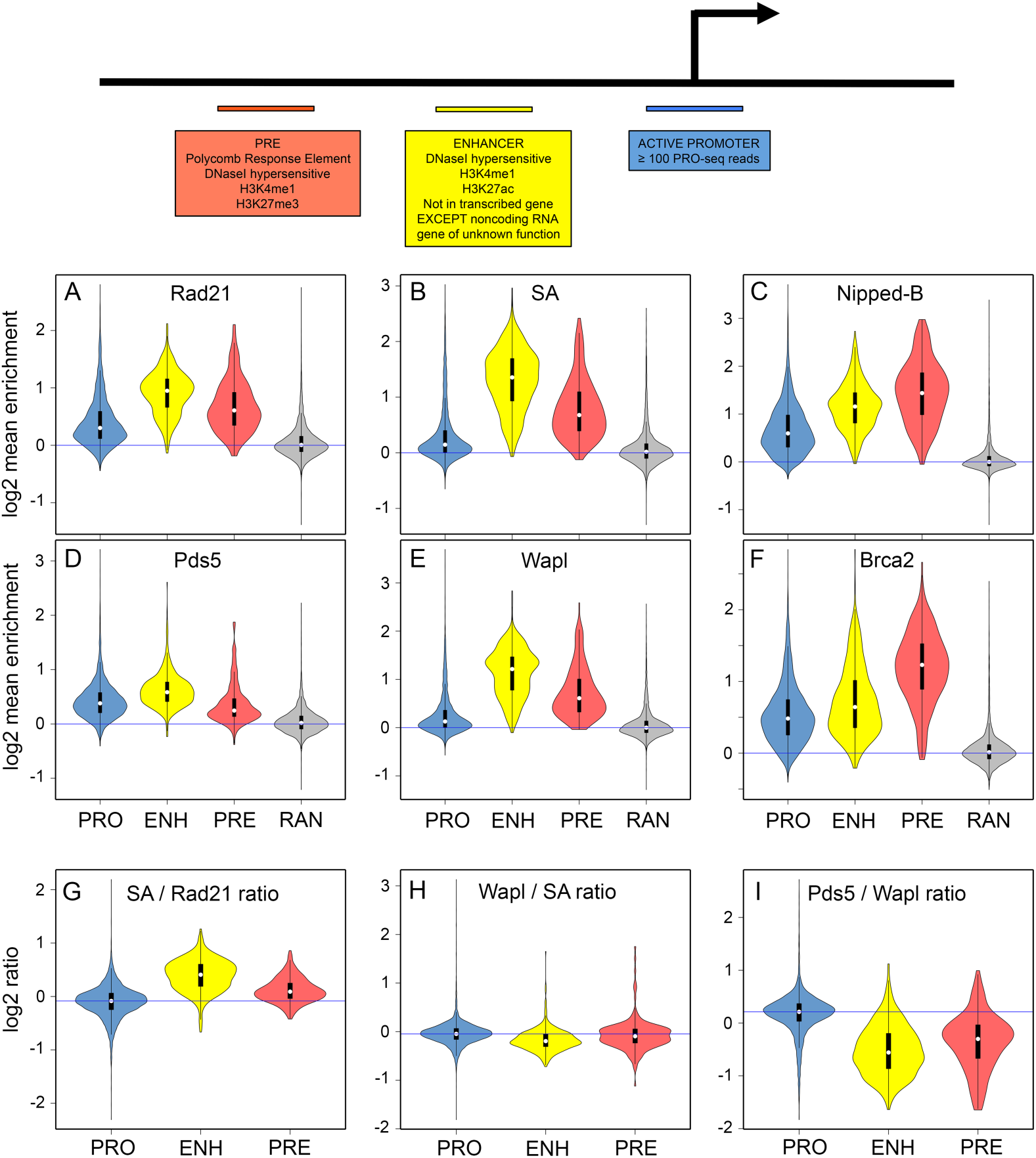
The relative levels of cohesin, Nipped-B, Pds5, Wapl and Brca2 vary between active promoters, enhancers and Polycomb Response Elements (PREs). The top diagram summarizes how active promoters (blue) extragenic enhancers (yellow) and PREs (orange) are defined as 500 bp sequences as described elsewhere [17, 33, 35]. There are 7,389 non-heterochromatic active promoters, 523 extragenic enhancers and 195 PREs. There are over 2,500 total active enhancers in BG3 cells but intragenic enhancers are excluded to avoid effects caused by changes in transcription. (**A**) Violin plots of the distribution of Rad21 ChIP-seq enrichment values (mean enrichment in each 500 bp element) for promoters (PRO, blue), extragenic enhancers (ENH, yellow) and PREs (PRE, orange), and 6,892 random 500 bp sequences as a negative control. White dots show the median values. (**B**) Same as A for SA. (**C**) Same as A for Nipped-B. (**D**) Same as A for Pds5. (**E**) Same as A for Wapl. (**F**) Same as A for Brca2. (**G**) Distribution of SA to Rad21 ChIP-seq enrichment ratios for promoters (PRO) enhancers (ENH) and PREs (PRE). (**H**) Same as G for the Wapl to SA ratio. (**I**) Same as G for the Pds5 to Wapl ratio.

The Rad21 and SA cohesin subunits show the highest levels at enhancers, followed by PREs, and promoters (Fig 6A,B). In contrast, Nipped-B has the highest levels at PREs, followed by enhancers and promoters (Fig 6C). Pds5 is highest at enhancers, followed by promoters, and PREs (Fig 6D) while Wapl shows the same pattern as cohesin, being higher at PREs than promoters (Fig 6E). Unexpectedly, the Brca2 pattern is more similar to Nipped-B than it is to Pds5, being highest at PREs, followed by enhancers and promoters (Fig 6F). The variation in the relative amounts of cohesin and accessory factors at different types of regulatory sequences imply that proteins that regulate transcription, and/or differences in chromatin structure at the regulatory sequences differentially influence their ability to bind cohesin and accessory factors. Each type of regulatory sequence also shows a relatively broad distribution of enrichments, which likely reflects differences in the proteins that bind to them in addition to their distance from an origin.

The varying ratios of cohesion factors between promoters, enhancers, and PREs suggest that the cohesin binding dynamics and function might differ between these types of gene regulatory sequences. For instance, the SA to Rad21 cohesin subunit ratio is highest at enhancers, followed by PREs and promoters while the Wapl to SA ratio is virtually identical at all regulatory sequences (Fig 6G,H). Thus cohesin composition, which could influence cohesin function, appears to differ between enhancers and promoters.

The effects of depleting cohesin accessory factors on cohesin levels and composition supports the idea that cohesin dynamics vary between different types of regulatory sequences. The Pds5 to Wapl ratio is substantially higher at promoters than enhancers or PREs (Fig 6I). Based on in vivo FRAP data showing that a high Pds5 to Wapl ratio destabilizes cohesin binding [9] the high Pds5 to Wapl ratio at promoters predicts that cohesin removal is more efficient at promoters than at enhancers. Supporting this idea, Pds5 depletion substantially increases Rad21 at promoters, with more moderate increases at PREs and very modest increases at enhancers (Fig 7A). All increases are statistically significant (S7 Table).

**Fig 7.**
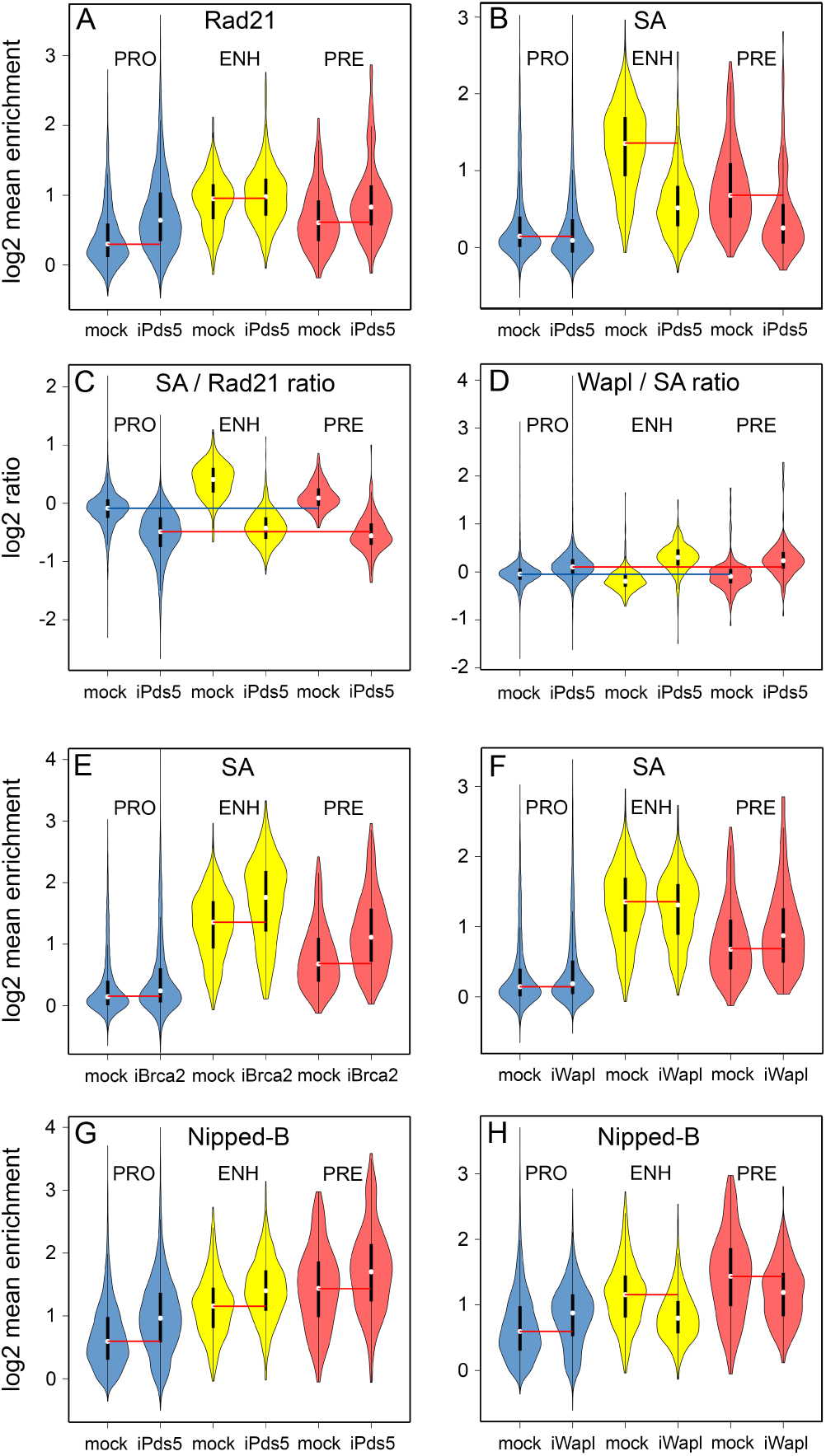
Pds5, Brca2 and Wapl differentially influence the levels of cohesin subunits and Nipped-B at gene regulatory sequences. (**A**) The violin plots show the distributions of Rad21 ChIP-seq enrichment values at promoters (PRO, blue) enhancers (ENH, yellow) and PREs (PRE, orange) in mock-treated control BG3 cells (mock) and cells depleted for Pds5 (iPds5). Red lines indicate the median values for each type of regulatory sequence in mock control cells. (**B**) Same as A for SA enrichment. (**C**) Distributions of the SA to Rad21 ratios at promoters (PRO) enhancers (ENH) and PREs (PRE) in control (mock) and Pds5-depleted (iPds5) BG3 cells. The blue line indicates the median ratio at promoters in mock control cells and the red line indicates the median ratio at promoters in Pds5-depleted cells. (**D**) Same as C for the Wapl to SA ratio. (**E**) Same as A for SA in control and Brca2-depleted (iBrca2) cells. (**F**) Same as A for SA in control and Wapl-depleted (iWapl) cells. (**G**) Same as A for Nipped-B enrichment. (**H**) Same as A for Nipped-B in mock control and Wapl-depleted (iWapl) cells. Statistical tests of the differences in the distributions of ChIP-seq enrichment after protein depletions in panels A, B, E, F, G and H are provided in S7 Table.

Although Pds5 depletion increases Rad21 levels, it also significantly lowers SA at all regulatory sequences (Fig 7B, S7 Table). The combined effect of increasing Rad21 and lowering SA equalizes the SA to Rad21 ratio at all regulatory sequences (Fig 7C). Pds5 depletion also slightly increases the Wapl to SA ratio at all regulatory sequences (Fig 7D). Brca2 depletion significantly increases SA levels at all regulatory sequences as expected from the metaorigin analysis above (Fig 7E, S7 Table). Wapl depletion slightly increases SA at promoters and PREs, with a minor statistically significant decrease at enhancers (Fig 7F, S7 Table). We hypothesize that the changes in cohesin levels and composition at gene regulatory sequences caused by depletion of Pds5, Wapl and Brca2 could alter cohesin dynamics and function at these sequences.

In addition to the multiple effects on cohesin, Pds5 and Wapl also influence binding of the Nipped-B cohesin loader at gene regulatory sequences. Pds5 depletion significantly increases Nipped-B at all regulatory sequences (Fig 7G, S7 Table) supporting the idea that Pds5 and Nipped-B compete for binding to cohesin. Wapl depletion substantially reduces Nipped-B at enhancers and PREs, but unexpectedly increases Nipped-B at promoters (Fig 7H, S7 Table). Wapl is low at promoters, so this suggests that regulatory sequences with high Wapl levels might sequester Nipped-B, and that proteins at promoters, potentially cohesin, or the TBPH RNA-binding protein [17] recruit Nipped-B released from enhancers and PREs by Wapl depletion. The idea that Wapl sequesters Nipped-B is consistent with the metaorigin analysis described above indicating that Wapl facilitates Nipped-B binding.

The interplay and variations between cohesin and the accessory proteins at gene regulatory sequences described above has important implications for how cohesin facilitates enhancer-promoter communication. It is widely postulated that cohesin holds enhancers and promoters together by an intra-chromosomal cohesion mechanism similar to how it holds sister chromatids together. However, the different SA to Rad21 ratios and levels of cohesin loading and removal factors at enhancers and promoters is inconsistent with this idea, which predicts that the cohesin populations at enhancers and promoters should be similar in composition and have similar dynamics. Instead, the SA to cohesin ratio is higher at enhancers than at promoters, and Pds5 depletion dramatically increases cohesin levels at promoters with little effect at enhancers, indicating that the cohesin populations at these sites are different complexes. We cannot rule out, however, the possibility that only a small fraction of cohesin molecules at enhancers and promoters participate in enhancer-promoter cohesion, and the majority of cohesin population detected by ChIP-seq do not.

### Pds5, Brca2 and Wapl have similar effects on gene expression as Nipped-B and cohesin

The finding that the levels of the different cohesin accessory factors and their ratios relative to cohesin vary between different types of regulatory sequences led us to hypothesize that their effects on gene expression would differ. Surprisingly, however, as described below, RNA-seq experiments reveal that despite differences in magnitude, the effects of Pds5, Brca2, and Wapl on gene expression significantly overlap those of Nipped-B and cohesin, largely affecting the same genes in the same directions. Our laboratory reported expression microarray and precision run-on sequencing (PRO-seq) experiments showing that Nipped-B and cohesin have very similar genome-wide effects on RNA accumulation and transcription in BG3 cells [14, 35]. For the studies here we performed RNA-seq in BG3 cells depleted for Nipped-B, Rad21, Pds5, Wapl, and Brca2, and compared their genome-wide effects on RNA levels (Fig 8). Three biological replicates were used for each depletion, and were compared to six mock control replicates. The Pearson correlation coefficients between replicates for genome-wide expression levels were all >0.95. The expression data are provided in S8 Data.

**Fig 8.**
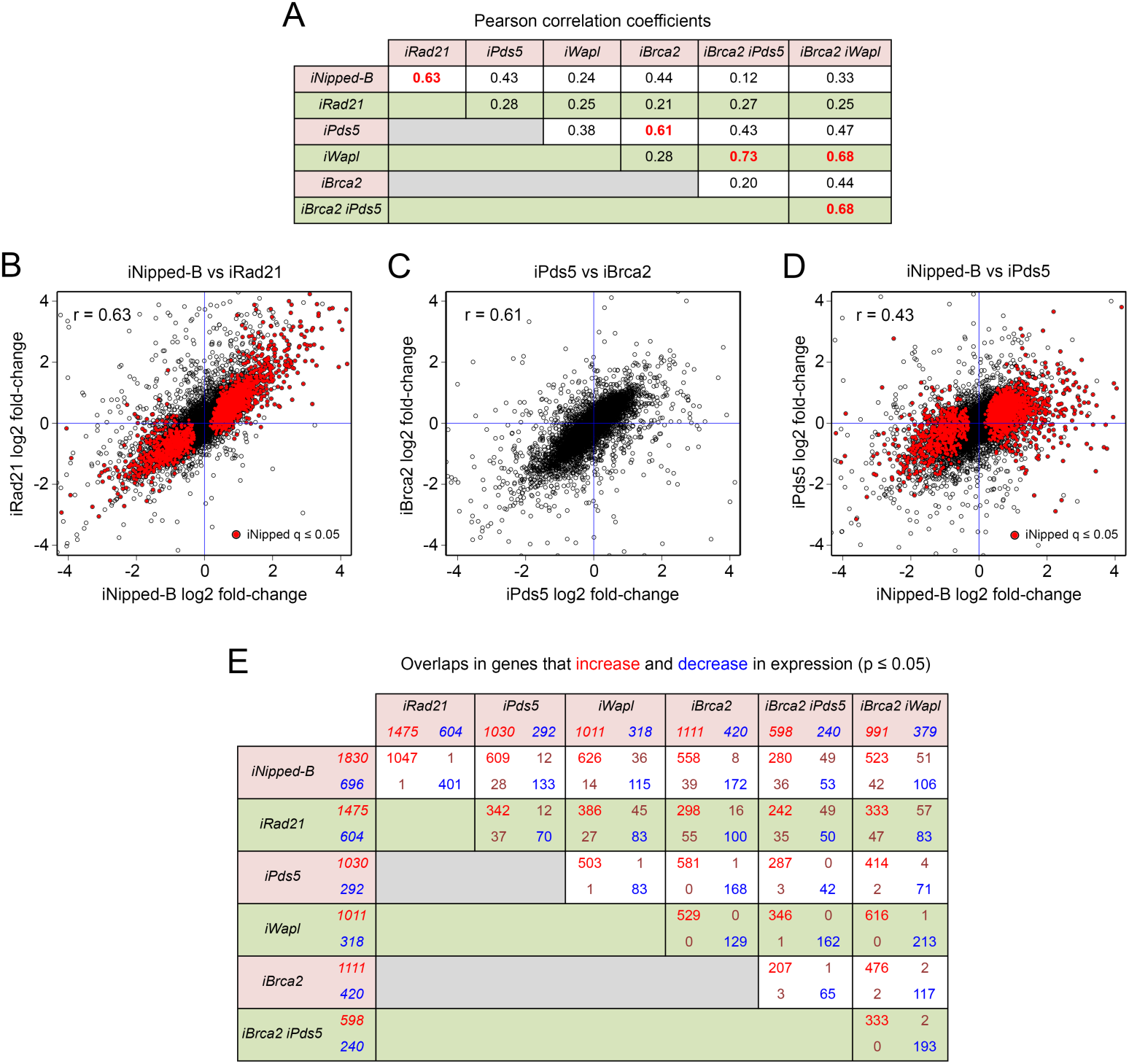
Pds5 and Brca2 have similar effects on gene expression in BG3 cells that overlap the effects of Nipped-B and cohesin. (**A**) Genome-wide Pearson correlation coefficients for the log2 fold-changes in RNA levels caused by depletion of Nipped-B (iNipped-B), Rad21 (iRad21), Pds5 (iPds5), Wapl (iWapl), Brca2 (iBrca2), Brca2 and Pds5 (iBrca2 iPds5) and Brca2 and Wapl (iBrca2 iWapl). Gene expression values used for the analysis are in S8 Data. (**B**) Dot plot of log2 fold-changes in RNA levels caused by Nipped-B depletion versus the changes caused by Rad21 depletion. Red dots show statistically significant changes in gene expression caused by Nipped-B depletion (q ≤ 0.05). (**C**) Dot plot of log2 fold-changes in RNA levels caused by Pds5 depletion versus changes caused by Brca2 depletion. (**D**) Dot plot of log2 fold-changes in RNA levels caused by Nipped-B depletion versus the changes caused by Pds5 depletion. Red dots show genes significantly altered by Nipped-B depletion (q ≤ 0.05). (**E**) Overlap in the genes that increase and decrease in expression with the indicated depletions at p ≤ 0.05. P values were used instead of the more stringent q values to obtain larger groups of genes. Numbers in red indicate genes that increase in expression and numbers in blue are genes that decrease. The numbers in the overlap boxes show the number that change with both depletion treatments. Red indicates genes that increase with both and blue indicate genes that decrease with both. Brown indicates genes that increase with one treatment, and decrease with the other. All overlaps in genes that increase or decrease in expression are statistically significant by Fisher’s exact test (S8 Data).

The genome-wide Pearson correlation between the fold-change changes in RNA levels caused by Nipped-B and Rad21 depletion is 0.63 (Fig 8A,B) consistent with prior studies showing that Nipped-B and cohesin have very similar effects on gene expression and transcription [14, 35]. The correlation between Pds5 and Brca2 depletion is 0.61, similar to the correlation between Nipped-B and Rad21 (Fig 8A,C). In contrast, the Pearson correlations between the effects of Nipped-B or Rad21 depletion versus Pds5 or Brca2 depletion are lower, ranging from 0.21 to 0.43 (Fig 8A,D). The correlations between Wapl depletion with Nipped-B, Rad21, Pds5 or Brca2 depletion range from 0.24 to 0.38 (Fig 8A).

The high correlation between the effects of Pds5 and Brca2 depletion on RNA levels was unexpected because Pds5 depletion reduces sister chromatid cohesion and Brca2 depletion does not. We thus conclude that most effects of Pds5 depletion on gene expression are not caused by sister chromatid cohesion defects. Because Pds5 depletion reduces cell division and Brca2 depletion does not, we also conclude that most changes in gene expression are not caused by changes in the cell cycle. It is possible, however, that some differences between the effects of Pds5 and Brca2 depletion reflect changes in sister chromatid cohesion or cell cycle effects.

The similar effects of Pds5 and Brca2 on gene expression make it unlikely that the effects on gene expression reflect a change in overall chromosome architecture because Pds5 depletion substantially alters cohesin distribution and Brca2 depletion does not. Also, if enhancer-promoter looping involves an intra-sister cohesion mechanism similar to the mechanism that holds sister chromatids together, then Pds5 depletion, which ablates sister chromatid cohesion, should have a different effect than Brca2 depletion. It is also possible, however, that most effects of Pds5 and Brca2 on gene expression do not involve changes in chromosome looping. Pds5 depletion reduces SA levels on promoters, enhancers and PREs, while Brca2 depletion increases SA levels. Thus the changes in gene expression caused by Pds5 or Brca2 depletion are also unlikely to be caused by changes in SA occupancy.

Although the correlations between the effects of Nipped-B or Rad21 depletion with those of Pds5, Brca2 and Wapl depletion are relatively low, close examination indicates that the same genes are affected. Most genes affected by Nipped-B depletion are altered in the same direction by Pds5 depletion as illustrated by the dot plot in Figure 8D, although the magnitudes of the changes tend to be smaller with Pds5 depletion. Agreeing with prior studies [14] gene ontology analysis shows that genes involved in neurogenesis and imaginal disc development are increased by all depletions, and that genes involved in protein translation are reduced (S8 Data). Moreover, there is significant overlap in the genes that increase in expression (p ≤ 0.05) with Nipped-B or Rad21 depletion and the genes that increase with Pds5, Wapl, or Brca2 depletion, as shown in Figure 8E. All overlaps in genes that increase with depletion of any of the cohesion factors are significant by Fisher’s exact test (S8 Data) and few genes show changes in the opposite directions. There is also statistically significant overlap in the genes that decrease with Nipped-B or Rad21 depletion and the genes that decrease with depletion of Pds5, Wapl, or Brca2, with only a few opposite effects, primarily with Wapl depletion (Fig 8E).

Although the effects of individually depleting Pds5 and Brca2 correlate modestly with Wapl depletion (0.38, 0.28) co-depletion of Pds5 and Brca2 strongly correlates with Wapl depletion (0.73) and Wapl-Brca2 co-depletion (0.68) (Fig 8A). The Pds5-Brca2 and Wapl-Brca2 double depletions show more modest correlations with Nipped-B or Rad21 depletion (0.12 – 0.33) or with depletion of Pds5 or Brca2 alone (0.20 –0.47) than with Wapl depletion. There is still significant overlap in the genes that increase in expression with Nipped-B or Rad21 depletion, but more genes that decrease in expression with Nipped-B or Rad21 depletion show increased expression in the double depletions (Fig 8E). Because Wapl depletion has virtually the same effects as Pds5-Brca2 or Wapl-Brca2 co-depletion, we conclude that loss of Wapl is epistatic to loss of Pds5 or Brca2, suggesting that much of the influence of Pds5 and Brca2 on gene expression requires Wapl. Wapl depletion more frequently has an opposite effect on those genes that decrease in expression upon Nipped-B or cohesin depletion, suggesting that Wapl counteracts activation by Nipped-B and cohesin at some genes.

### Brca2 influences gene expression in developing wings and facilitates growth

The finding that Brca2 influences gene expression in BG3 cells raised the question of whether or not it also influences gene expression during in vivo development. We conducted RNA-seq in 3^rd^ instar wing imaginal discs from different control genetic backgrounds, and two different *brca2* null mutants, which revealed that many genes increase and decrease in expression. Using a statistical threshold of q ≤ 0.05 (5% false discovery rate) 208 genes increase in expression and 606 decrease, with most changes less than 2-fold (Fig 9A, S8 Data). Gene ontology analysis indicates that the decreasing genes are involved in morphogenesis and development (S8 Data). In additional to many genes that regulate development, the genes that decrease in expression include broadly-acting transcription factors such as Mediator subunits and Pol II kinases, cell cycle control genes, and genes encoding DNA repair proteins (Fig 9B). Unlike *pds5* or *wapl* null mutant flies [7, 8] *brca2* null mutant flies are viable, although females are sterile because of defective meiosis [55].

**Fig 9.**
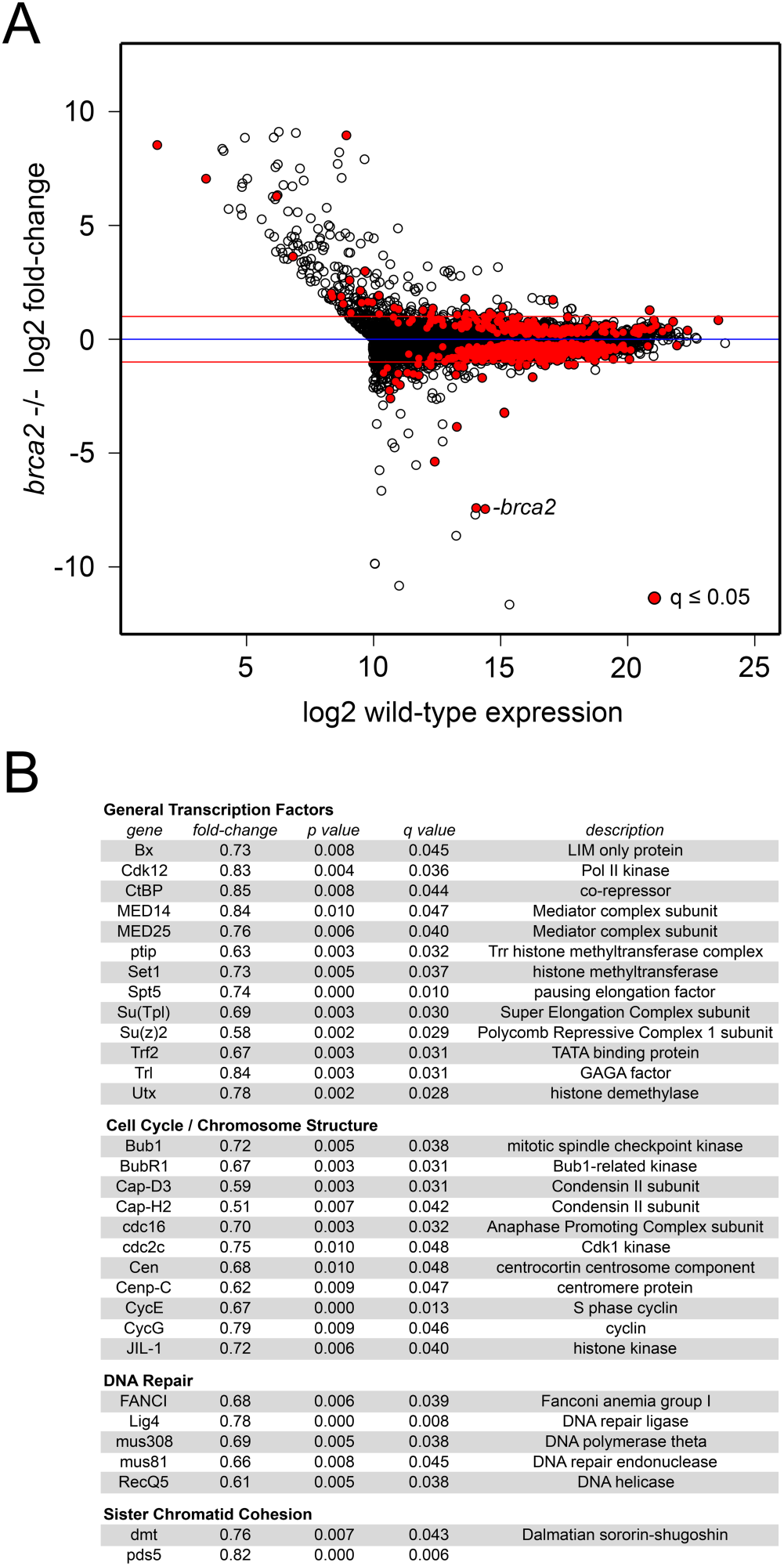
Brca2 influences gene expression in developing wings. (**A**) The log2 fold-change in gene expression in *brca2* null mutant 3^rd^ instar wing imaginal discs is plotted versus the log2 expression level in control wing discs for all active genes. Active genes are defined as those that are expressed at or above the median level in control discs plus those expressed at or above the control median level in *brca2* mutant discs. Red dots indicate statistically significant changes in gene expression (q ≤ 0.05). The dot representing the *brca2* gene is labeled. The blue line indicates no change, and the two red lines indicate 2-fold increases or decreases. (**B**) Examples of genes down-regulated in *brca2* mutant discs in the indicated categories. This is not a comprehensive list, which can be generated from the expression data provided in S8 Data.

There are no overt structural mutant phenotypes in *brca2* null adult flies, but it seems likely that the many modest changes in gene expression could cause subtle growth or developmental deficits that might be revealed by close examination, or in different genetic backgrounds. We measured the sizes of adult male and female *brca2* mutant wings, to look for possible changes in growth. Male *brca2* mutant wings are some 9% smaller than controls, and female wings are 4% smaller (S9 Fig). These reductions are similar in magnitude to the dominant effects of null mutations in the fly *myc* (*diminutive*, *dm*) and *Tor* genes encoding critical dosage-sensitive growth regulators [56]. They are also similar to the effects of heterozygous *Nipped-B* mutations [56]. Thus Brca2 facilitates wing growth, and we hypothesize that this reflects the role of Brca2 in gene expression.

## Discussion

The studies presented here show that the Pds5-Wapl complex limits the size of the cohesin binding domains centered around DNA replication origins, and that Pds5 and Brca2, which form a complex lacking Wapl [10–12] have opposing effects on sister chromatid cohesion and binding of the SA cohesin subunit near early replication origins. In contrast to their opposing roles in sister cohesion, Pds5 and Brca2 have very similar influence on gene expression, affecting largely the same genes as Nipped-B and cohesin. As outlined below, these findings have significant implications for where cohesin binds, and cohesin’s roles in sister chromatid cohesion and gene regulation. Some of the key hypotheses are outlined in Figure 10. Our observations make predictions about mutations that could cause human developmental syndromes, and have implications for how *BRCA2* mutations might increase cancer susceptibility.

**Fig 10.**
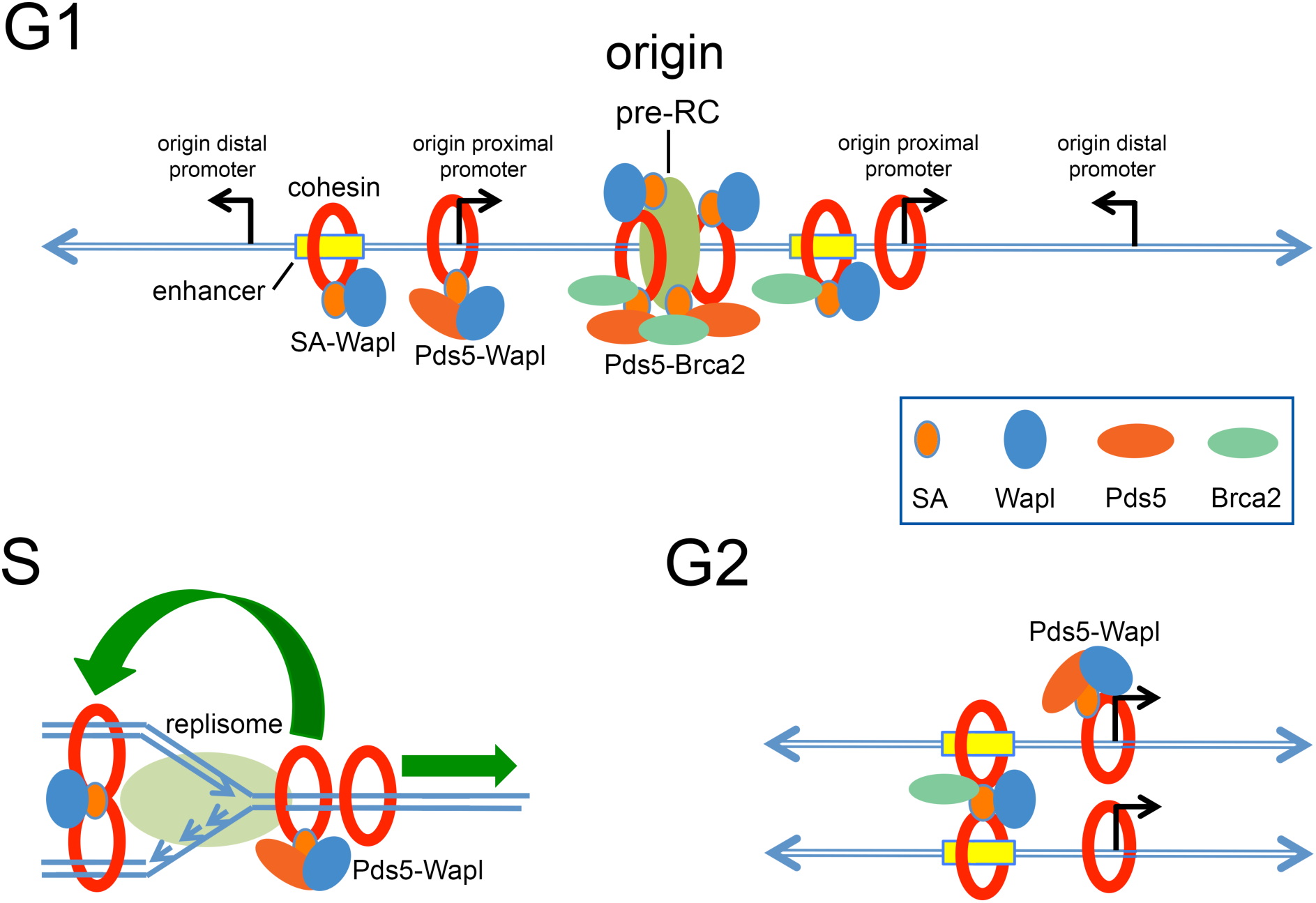
Models for the roles of Pds5, Wapl and Brca2 in regulating cohesin dynamics and function in Drosophila cells. At the top, in the G1 phase of the cell cycle, active promoters (angled arrows) and enhancers (yellow boxes) proximal to DNA replication origins bind cohesin (red rings) while origin-distal promoters and inactive genes do not. For simplicity, the Nipped-B cohesin-loading factor is not depicted, but it is present wherever there is cohesin. We posit that book-marking proteins (not depicted) at promoters and enhancers remain bound through mitosis to recruit Nipped-B and enable cohesin loading after cell division. Enhancers have a relatively high level of the SA cohesin subunit (small orange oval) compared to promoters. Wapl (blue oval) is stoichiometric with SA, and unlike Pds5 (large orange oval) binds everywhere with cohesin. Promoters have a high Pds5 to Wapl ratio, and the Pds5-Wapl cohesin removal complex keeps cohesin levels comparatively low. The pre-replication complex (large light green oval) containing the origin recognition complex (ORC) and the MCM helicase complex licenses DNA replication origins (origin) in early G1 and recruits Nipped-B resulting in cohesin loading and binding of Pds5, Wapl, and Brca2. Pds5 aids and Brca2 inhibits SA binding at origins to titrate the fraction of cohesin complexes that mediate sister cohesion during G2. At origins, Pds5 and Wapl do not remove cohesin. Pds5 restricts binding of Nipped-B via competition for cohesin. During S phase (lower left) the replisome pushes cohesin ahead of the replication fork. The Pds5-Wapl complex unloads cohesin in front of the fork, limiting cohesin spreading. Cohesin is reloaded behind the fork to establish sister chromatid cohesion, which requires SA. A handcuff model is shown, but cohesion mechanisms with single cohesin rings are possible. During G2 (lower right) enhancers have a high SA to cohesin ratio, and a low Pds5 to Wapl ratio, which may indicate that they are sites of sister chromatid cohesion. Cohesion is low at promoters, which have a low SA to cohesin ratio, and a high Pds5-Wapl ratio. We speculate that unrestrained promoters loop independently to a double enhancer complex held together by sister cohesion, aided by interactions between Nipped-B, cohesin and the Mediator complex (not depicted). Pds5, Brca2 and Wapl control cohesin binding dynamics and cohesin-dependent looping to influence gene expression.

### Cohesin localization

The finding that the maximal levels of cohesin and cohesin regulatory factors center at replication origins and decrease extending outward for many kilobases strongly suggests that DNA replication plays a role in positioning cohesin. This hypothesis is consistent with single molecule studies in Xenopus extracts showing that DNA replication causes cohesin to translocate unidirectionally [41]. The single-molecule studies also show that cohesin translocation is suppressed by Pds5 and Wapl [41]. This agrees with our finding that depletion of Pds5 or Wapl increases cohesin levels for tens of kilobases surrounding replication origins. We thus hypothesize that pushing of cohesin by replication forks is a key determinant of cohesin localization and which genes bind cohesin, and that reduction of Pds5 or Wapl slows the rate of cohesin removal in front of replication forks, leading to increased cohesin domain size (Fig 10).

Alternatively, it is possible that the bulk of cohesin loading occurs at licensed origins early in G1, and that cohesin diffuses bidirectionally to be trapped at active genes and regulatory sequences. As active genes trap cohesin, there would be fewer cohesin rings to diffuse further from the origin. In this scenario, reducing the rate of cohesin removal by depleting Pds5 or Wapl increases the number of topologically-bound cohesin rings available to translocate past origin-proximal genes. It will require new and precise synchronization methods to obtain cells in the appropriate cell cycle stages (early G1, late G1, early S) to distinguish between these possibilities. We currently favor the idea that DNA replication pushes cohesin based primarily on the in vitro experiments showing that replication forks push cohesin [41] and yeast studies indicating that cohesin can travel with replication forks [32].

In Xenopus oocyte extracts, association of the Nipped-B orthologs with chromatin and cohesin loading require replication origin licensing and formation of the complete pre-replication complex (pre-RC) containing the origin recognition complex (ORC) the minichromosome maintenance (MCM) helicase complex, Cdc6 (cell division cycle 6) and Cdt1 (cyclin-dependent transcript 1) [27–30]. This supports the idea that replication origins are major loading sites for cohesin. Cohesin is also enriched at origins in HeLa cells [31]. However, studies in Drosophila cultured cells depleted for pre-RC components indicate that origin licensing is not essential for stable association of cohesin with chromatin, although these studies did not examine cohesin localization [26]. Xenopus oocytes do not have active gene transcription, and we think it likely that a combination of both licensed replication origins and active genes determine cohesin binding in Drosophila cells (Fig 10). Depletion of pre-RC components blocks DNA replication, and thus precise cell synchronization and rapid protein degradation methods will be needed to fully dissect the roles of replication origins, DNA replication and active genes in cohesin positioning.

With these caveats in mind, we currently envision that DNA replication exerts global control over the extent of cohesin domains, and that within these domains, proteins binding to active genes and regulatory sequences define the detailed cohesin localization and dynamics. Once an active gene or regulatory sequence binds cohesin and accessory factors, cohesin is continuously loaded and removed from these sites during interphase [9]. In this scenario, the cohesin dynamics at different sites is fine-tuned by the ratios of Nipped-B, Pds5, Wapl and Brca2, which as shown here, vary between promoters, enhancers and PREs. A key question this idea raises, however, is how cohesin is positioned on chromosomes in early G1 after mitosis and before DNA replication. We speculate that book-marking factors remain bound to cohesin-binding genes and regulatory elements through mitosis and determine cohesin re-loading in early G1 together with the proteins binding active genes and regulatory elements. The cohesin and accessory protein levels for each gene and regulatory sequence are then refreshed during DNA replication as necessitated by duplication of the genome. This model predicts that the putative mitotic bookmarks may also be extended to additional active genes upon Pds5 or Wapl depletion as the cohesin domains increase in size.

Studies in yeast implicate DNA replication in temporarily positioning cohesin [32] and cohesin is enriched at origins in HeLa cells [31] but a potential role for DNA replication in cohesin localization in mammalian cells remains to be investigated. Recent studies show that transcription, the CTCF architectural protein, and Wapl position cohesin in mammalian cells [57]. Absence of CTCF causes cohesin to localize to transcription start sites, similar to the pattern observed in Drosophila. Unlike mammalian CTCF, Drosophila CTCF does not interact directly with the SA cohesin subunit [58] and cohesin does not significantly co-localize with CTCF [36]. Thus the pattern observed in CTCF-deficient mammalian cells is similar to the normal pattern in Drosophila.

In mammalian cells that lack both CTCF and Wapl, cohesin accumulates in large islands several kilobases in size at the 3’ ends of genes [57] suggesting that transcription pushes cohesin to the ends of genes, as seen in yeast [59–61]. Although we detect modest increases in cohesin in some intergenic regions upon Pds5 or Wapl depletion in Drosophila cells, we do not see new substantial intergenic domains, suggesting that there may be additional factors in mammalian cells that trap cohesin at the 3’ ends of genes. Although it remains to be investigated, we envision that DNA replication also positions cohesin in mammalian cells, and that CTCF and transcription refine cohesin positioning, similar to our finding that the cohesin extending out from Drosophila replication origins skips over inactive genes to associate with active genes and regulatory sequences.

### Sister chromatid cohesion

It is poorly understood why Pds5, which removes cohesin from chromosomes, is also required for sister chromatid cohesion [62]. A recent study in budding yeast intimately link the role of Pds5 in cohesion to DNA replication by showing that mutations affecting the Elg1 protein that unloads the PCNA replication clamp, or alternatively overexpression of PCNA, suppress cohesion defects of *pds5* mutants [63]. Our findings lead us to speculate that Pds5 activity is modified at replication origins, where it actively increases the levels of the SA subunit, and that the high level of SA at origins is important for establishing sister cohesion during replication. SA is required for sister cohesion although it is not part of the ring that encircles DNA, and cohesin topologically binds chromosomes without SA [43]. It is proposed that SA can link two cohesin rings to mediate a handcuff mechanism for sister chromatid cohesion [44, 45] or alternatively that SA may act as a chaperone that facilitates topological binding of one cohesin ring around both sisters at replication forks [43]. These reports, together with the correlation between the loss of high SA levels at replication origins and loss of sister chromatid cohesion upon Pds5 depletion, lead us to propose that maintaining high levels of SA at origins is the critical Pds5 function for establishing sister chromatid cohesion. This idea is further supported by the finding that Brca2 depletion suppresses both the loss of cohesion and the loss of SA at origins caused by Pds5 depletion. Wapl depletion does not alter SA levels at origins, and does not reduce sister chromatid cohesion. An origin-specific role for Pds5 in facilitating SA binding can thus resolve the paradox that Pds5 is required for both sister chromatid cohesion and cohesin removal.

We do not know how Pds5 facilitates SA association with origins or how Brca2 counteracts this activity. We can envision, however, multiple possible direct mechanisms. SA, like Pds5, Wapl and Nipped-B contains several HEAT armadillo-like repeats, and has a structure similar to these cohesin regulators [13, 22, 46–50, 64, 65]. We thus speculate that when brought into proximity of each other on chromosomes, Brca2 might also interact with SA, or even Nipped-B. Although Wapl did not co-purify with Brca2 from Drosophila extracts, significant sub-stoichiometric amounts of SA and Nipped-B were detected [11]. Pds5 and Nipped-B proteins form similar hook-like structures and interact with multiple cohesin subunits. Brca2 interacts with a region in the “handle” of Pds5 adjacent to the N terminal region that interacts with Wapl and facilitates Pds5 dimerization [11, 13]. Thus one idea is that Pds5 aids SA binding through interactions with Rad21 and SA, but Pds5 dimers formed by Brca2 do not. Brca2 depletion would reduce Pds5 dimerization and thereby increase SA binding. The caveat to this argument, however, is that Brca2 depletion also reduces Pds5 binding, and thus should also reduce SA binding. An alternative idea is that Brca2 might interact with SA on chromosomes and reduce its affinity for cohesin. It is also possible that the pre-RC at origins facilitates SA binding, and that Brca2 inhibits this binding via interaction with the pre-RC or SA. In this case, Pds5 could facilitate SA loading by interacting with Brca2 and preventing it from interacting with SA. Testing these ideas may be possible using Xenopus extracts that couple origin licensing with cohesin loading [27–30] and by examining the direct interactions of SA, Pds5 and Brca2 with each other and with replication factors.

As described below, although Pds5 and Brca2 have opposing effects on sister chromatid cohesion and SA levels at replication origins, they have very similar effects on gene expression. This apparent paradox can be explained if Pds5 activity is altered at origins or in early G1 during origin licensing. In other words, we suspect that the roles of Pds5 in sister chromatid cohesion and gene expression are separated by chromosomal location and/or phase of the cell cycle.

The Pds5-Brca2 complex is important for mitotic DNA repair and homologous recombination during meiosis, corresponding with the known role of Brca2 in DNA repair [10–12]. The finding that Brca2 can oppose Pds5 function in sister chromatid cohesion shows that Brca2 plays an additional role in regulating genome stability even in the absence of DNA damage. Although increased SA at origins is apparent with Brca2 depletion alone, *brca2* null mutant flies are viable, and the anti-cohesion effects of Brca2 are not apparent unless Pds5 is reduced. The anti-cohesion role of Brca2 is also opposite to its role in DNA repair in the sense that reduced cohesion decreases genome stability and DNA repair increases stability. It remains to be seen if hypermorphic *brca2* mutations or overexpression of Brca2 can reduce sister chromatid cohesion sufficiently to alter chromosome segregation, but this is potentially relevant to the increased cancer susceptibility in some individuals with *BRCA2* missense mutations. It is also of interest that the human *BRCA2* gene neighbors *PDS5B*, with some cancer-associated mutations likely altering both genes [66]. This raises the possibility that some mutations could alter the BRCA2-PDS5B ratio in a way that disfavors sister chromatid cohesion.

### Gene regulation

Since the original discovery that *Nipped-B* mutations alter enhancer-dependent gene expression [67] an attractive concept has been that cohesin holds enhancers and promoters together by a topological mechanism similar to how it holds sister chromatids together. This idea has been expanded in loop extrusion models in which chromatin is threaded through topologically bound cohesin to form intra-chromosomal loops important for topological domain (TAD) formation and gene regulation [68]. These models thus also suggest cohesion-like mechanisms to hold together two regions of the same chromosome.

Not all TADs in Drosophila cells, however, require cohesin for their formation [36, 38–40] and data presented here argue against the intra-chromosomal cohesion model for facilitating enhancer-promoter loops. The Pds5 to Wapl ratio is substantially higher at promoters than at enhancers, which based on in vivo FRAP [9] predicts that cohesin binding is less stable at promoters (Fig 10). Indeed, as shown here, Pds5 depletion substantially increases cohesin levels at promoters but has much less effect at enhancers. The intra-chromosomal cohesion model predicts that the cohesin dynamics will be same at enhancers and promoters, either because they are the same cohesin complexes at both sequences in the embrace cohesion model, or are tightly linked to each other in the handcuff model. A caveat to this argument is that if there are several cohesin rings at these regulatory sequences, as few as one may mediate intra-chromosomal cohesion, while the majority differ in their composition and binding dynamics.

The finding that Pds5 and Brca2 depletion have very similar genome-wide effects on gene expression provides further evidence against the intra-chromosomal cohesion model because Pds5 depletion strongly reduces sister cohesion and Brca2 depletion does not. If sister chromatid cohesion is lost, then intra-chromosomal looping should also be substantially reduced. It is also possible, however, that only a small fraction of the effects of Pds5 depletion on gene expression involve looping deficits, and that changes in cohesin’s other roles in gene regulation are responsible for most effects on gene expression. For instance, cohesin influences transcription of active genes by recruiting the PRC1 Polycomb repressive complex to the promoter region [33, 53]. Nipped-B and cohesin bind to essentially all enhancers, however, and there are many instances where Nipped-B or cohesin depletion reduces transcription of genes activated by known enhancers [35]. We thus currently prefer the alternative idea that interactions between Nipped-B or cohesin with other proteins, such as the Mediator complex [69] facilitate enhancer-promoter looping.

The finding that Pds5 and Brca2 influence expression of the same genes as Nipped-B and cohesin supports the idea that they mediate their effects on gene expression in large part through their effects on cohesin binding and dynamics. It is counterintuitive, however, that their effects are largely in the same direction as Nipped-B or cohesin because Pds5 depletion increases cohesin at promoters and Nipped-B at all regulatory sequences. It is also unexpected that Pds5 and Brca2 depletion, which have opposite effects on SA levels at regulatory sequences, have the same effect on gene expression. These findings thus argue that optimal cohesin dynamics are more important than absolute cohesin levels for gene expression. It must be kept in mind that most genes that bind cohesin do not show large changes in transcription upon cohesin depletion, and that only a subset show substantial and statistically significant changes [14, 35]. Both increases or decreases in cohesin levels or dynamics at these more sensitive genes could have similar effects on their transcription.

Another possibility we considered is that the level of Pds5, and not Nipped-B or cohesin, is more crucial for gene regulation. Nipped-B and Brca2 depletion reduce Pds5 at all regulatory sequences except PREs, and thus have similar effects on Pds5 binding levels as depletion of Pds5. Against this idea, Wapl depletion increases Pds5 levels on gene regulatory elements, and has similar effects on gene expression as Pds5 depletion. We thus currently favor the idea that cohesin binding dynamics is the key factor that drives the majority of the significant effects of cohesin on transcription. This idea can be tested in future experiments through more direct measurements of transcription such as nascent RNA sequencing, and overexpression of cohesin regulators.

The finding that Brca2 has significant effects on gene expression in both cultured cells and in vivo has implications for the role of *BRCA2* mutations in cancer susceptibility. Many of the genes whose expression is altered in *brca2* mutant wing discs are involved in development, cell cycle control and DNA repair, or are broadly active transcriptional regulators. Thus effects on gene expression caused by *BRCA2* mutations could conceivably contribute to cancer etiology even in the absence of DNA damage.

### Can *PDS5A*, *PDS5B*, *WAPL* and *BRCA2* mutations cause cohesinopathy-like syndromes?

Heterozygous loss-of-function mutations in *NIPBL*, the human Nipped-B ortholog cause Cornelia de Lange syndrome (CdLS) which displays significant deficits in physical and intellectual growth and development, and structural abnormalities in the face, limbs and organs [70, 71]. Typically milder forms of CdLS are caused by dominant missense mutations in the *SMC1A* and *SMC3* cohesin subunit genes [72, 73]. Dominant loss of function mutations in *HDAC8* encoding the deacetylase that recycles acetylated SMC3 cause CdLS similar in severity to *NIPBL* mutations [74] and deficiencies in *RAD21* cause a developmental syndrome with deficits overlapping CdLS [75]. Individuals with cohesinopathy mutations do not display overt sister chromatid cohesion or chromosome segregation phenotypes, and thus the leading idea is that the developmental deficits arise from changes in gene expression [76, 77]. The finding that Pds5, Brca2 and Wapl depletion have similar effects on gene expression as Nipped-B and cohesin in BG3 cells, and that *brca2* null mutant wing discs show significant changes in the expression of many genes that influence growth and development raises the question of whether the human orthologs could also be cohesinopathy genes.

There are two Pds5 orthologs in mammals, and mice homozygous mutant for *Pds5A* or *Pds5B* show severe lethal developmental phenotypes [51, 78, 79] overlapping but not identical to those caused by heterozygous *Nipbl* loss-of-function mutations [80]. The individual *Pds5A* and *Pds5B* heterozygotes do not display overt phenotypes. It can be predicted, therefore, that similar mutations in humans would not cause a dominant syndrome, but would be recessive lethal.

Drosophila *wapl* hemizygous null mutations are lethal, but there are no overt adult phenotypes in heterozygous females, despite a measurable effect on global cohesin chromosome-binding dynamics [9]. Strikingly, however, a dominant-negative *wapl* mutant allele that produces a truncated Wapl protein stabilizes and increases cohesin binding and causes a Polycomb mutant phenotype reflecting decreased epigenetic silencing of homeotic genes [81]. *Nipped-B* and cohesin mutations dominantly suppress the dominant transformation phenotypes of *Pc* mutants [53, 82, 83] and thus it is possible that a *WAPL* dominant-negative mutation could cause a human developmental syndrome that would differ significantly from CdLS and other cohesinopathies, and which may be lethal in utero.

Although Drosophila *brca2* null mutants are viable [55] mouse *Brca2* null mutations are homozygous lethal during embryogenesis [84] and thus are also likely early recessive lethal in humans. However, some mice with *Brca2* hypomorphic mutations survive and are smaller than littermates, with multiple developmental abnormalities [85]. Importantly, biallelic human *BRCA2* mutations expressing truncated or otherwise altered BRCA2 proteins cause Fanconi anemia type D1 [86, 87]. This type of Fanconi anemia is associated with highly penetrant and fatal early childhood leukemia, but also multiple congenital abnormalities including slow growth and microcephaly, which are phenotypes that also occur in CdLS. The sources of the developmental deficits in *Brca2* mutant mice and Fanconi anemia D1 are unknown, but our results showing effects on gene expression and growth in developing Drosophila wings raise the likelihood that they reflect multiple changes in gene expression, similar to those caused by heterozygous *Nipped-B* and *NIPBL* mutations.

## Materials and methods

### Cell culture, RNAi treatment and metaphase spreads

BG3 cells were cultured and RNAi treatment and metaphase spreads were performed as previously described [14]. At least 50 metaphase nuclei were scored for each treatment group and time point in each experiment.

### Pds5 and Brca2 antibodies

Full length Pds5 (residues 1 – 1218) was expressed as a His6 fusion in *E. coli*, purified by nickel chromatography, and used to immunize a guinea pig at Pocono Rabbit Farm and Laboratory, Inc. Brca2 residues 1 to 404 of were expressed as a His6 fusion peptide in *E. coli*, purified and used to immunize a rabbit and a guinea pig at Josman, LLC.

### ChIP-seq

ChIP-seq was performed as described before [17, 18]. The Nipped-B, Rad21, SA and Wapl antibodies and their validation were described previously [34, 81]. The Pds5 and Brca2 antibodies were validated by RNAi westerns (S1 Fig). Multiple independent biological replicate experiments were performed for each experimental ChIP-seq group, sequencing to at least 10X genome coverage per replicate, and normalizing to input chromatin sequencing of at least 45X genome coverage. Mean enrichment at active promoters, enhancers and PREs was calculated as described before [17, 33] using custom R [88] scripts. Meta-origin analysis was performed using custom R scripts and the locations of early DNA replication origins in BG3 cells (GEO Accession GSE17281, S3 File). Meta-origin plots were made using Microsoft Excel. Violin plots were made using the wvioplot.R package (<u>https://github.com/cran/wvioplot</u>). The Integrated Genome Browser (IGB) [89] was used to inspect ChIP-seq data and prepare figures.

### DNA fiber analysis

DNA fibers were spread on microscope slides by adapting a published procedure [90]. Mock or RNAi-depleted BG3 cells were labeled with 20 micromolar IdU (ThermoFisher, cat no. 10035701) in the media for 20 min at 25°, washed twice with phosphate buffered saline (PBS) and then labeled for 20 min with 200 micromolar CldU in the culture media (Sigma Aldrich, cat no. C6891) at 25°. Labeled cells were washed twice with PBS, trypsinized, washed twice with PBS and brought to 7,000 cells per microliter in PBS and placed on ice. For experiments with hydroxyurea (HU) treatment, cells were washed twice with PBS after CldU incorporation and incubated at 25° with 4 mM HU for 4 hours prior to cell collection. Labeled cells were diluted with unlabeled control BG3 cells in a 1:5 or 1:10 ratio before spreading to decrease the number of overlapping labeled fibers [60]. Two microliters of diluted cells were mixed with eight microliters of lysis buffer (200 mM Tris·HCl pH 7.5, 50 mM EDTA, 0.5% SDS) on top of a microscope slide (VWR, cat no. 48311-703) for eight min. After lysis, the slides were tilted at a 20 to 40° angle to allow the DNA to spread. Slides were air-dried at room temperature for 1 hour and fixed in 3:1 v/v methanol:acetic acid solution for eight min. Fixed slides were dried and stored at 4° before immunostaining.

For immunostaining, fixed slides were washed twice for five min with PBS and treated with 2.5 N HCl for one hour. Acid-treated slides were washed three times for five min with PBS, blocked with 5% (w/v) bovine serum albumin (BSA) in PBS for one hour at 37° and rinsed in PBST (0.05% Tween 20 in PBS) for five to ten sec just prior to adding primary antibodies diluted in PBST containing 1% BSA. The primary antibodies used were 1:20 mouse anti-BrdU B44 (BD Biosciences, cat no. 347580) which recognizes IdU and 1:100 rat anti-BrdU monoclonal antibody BU1/75 (ThermoFisher, cat no. MA1-82088) which recognizes CldU. Forty microliters of the primary antibody mix was added to each slide, covered with a coverslip, and the slides were incubated in a dark humid chamber for 1.5 hours at room temperature. Coverslips were removed in PBS, and the slides washed three times for five min with PBST and kept in PBS until addition of the secondary antibodies. The secondary antibodies were each diluted 1:125 in PBST containing 1% BSA. The secondary antibodies used were Alexa Fluor 546 goat anti-mouse IgG1 (Invitrogen, cat no. A21123) and Alexa Fluor 488 chicken anti-rat IgG (Invitrogen, cat no. A21470). Forty microliters of diluted secondary antibody mix was added to each slide, covered with a coverslip, and the slides were incubated for one hour in a dark humid chamber at room temperature. Coverslips were removed in PBS, and the slides washed three times for five min in PBST and left in PBS before mounting. For mounting, slides were air-dried at room temperature in the dark, and mounted with 20 microliters of ProLong Gold antifade reagent (Invitrogen, cat no. P36930) under a coverslip. Slides were dried at room temperature in the dark and stored at 4° before imaging. Fluorescent micrographs were digitally captured using a Leica SP5 laser scanning confocal microscope with a 60x objective, and the lengths of connected red and green fibers were measured using NIH ImageJ software. Statistical analysis was conducted using R and violin plots generated using wvioplot.R.

### RNA-seq

Total RNA-seq seq with BG3 cells was performed using ribosomal depletion and processed as described previously [33, 56] using three independent biological replicates for each depletion and six previously reported mock control samples [33] (GEO Accession GSE100548). Genome-wide expression values between replicates showed correlations >0.95. Gene expression data in mean normalized nucleotide coverage per gene, and gene ontology analysis is provided in S8 Data. Total RNA-seq with dissected late third instar wing discs from female larvae was conducted as described before using ribosomal depletion [56]. To minimize genetic background effects, five wild-type controls consisted of two Canton S samples and three *y w* samples from independent stocks. The five *brca2 -/-* replicates consisted of two isolations of GFP negative larvae from a *brca2*^*KO*^ */ CyO, Kr-GFP* stock and three isolations from GFP negative larvae from a *brca2*^*56e*^ */ CyO, Kr-GFP* stock. All replicates gave genome-wide correlation values >0.95. Both *brca2* null alleles were the gift of Trudi Schüpbach. The expression data in mean normalized nucleotide coverage per gene, and gene ontology analysis is provided in S8 Data.

### Wing area measurements

Wings were collected from adult flies from crosses conducted at 25°, mounted, photographed and measured as previously described [56].

## Acknowledgements

We thank Kathie Mihindukulasuriya for construction of some RNA-seq libraries and operation of the DNA sequencer, Judy Kassis for providing Wapl antibody, Trudi Schüpbach for the *brca2*^*KO*^ and *brca2*^*56e*^ alleles, David MacAlpine for the processed BG3 early DNA replication data, the Alessandro Vindigni laboratory for advice on DNA fiber analysis, and Michael Rauchman for comments on the manuscript. This work was supported by grants from the National Institutes of Health (R01 GM108714) and the Alvin J Siteman Cancer Center, Washington University School of Medicine (8074-88) to DD.

## Supporting information

**S1 Fig.**
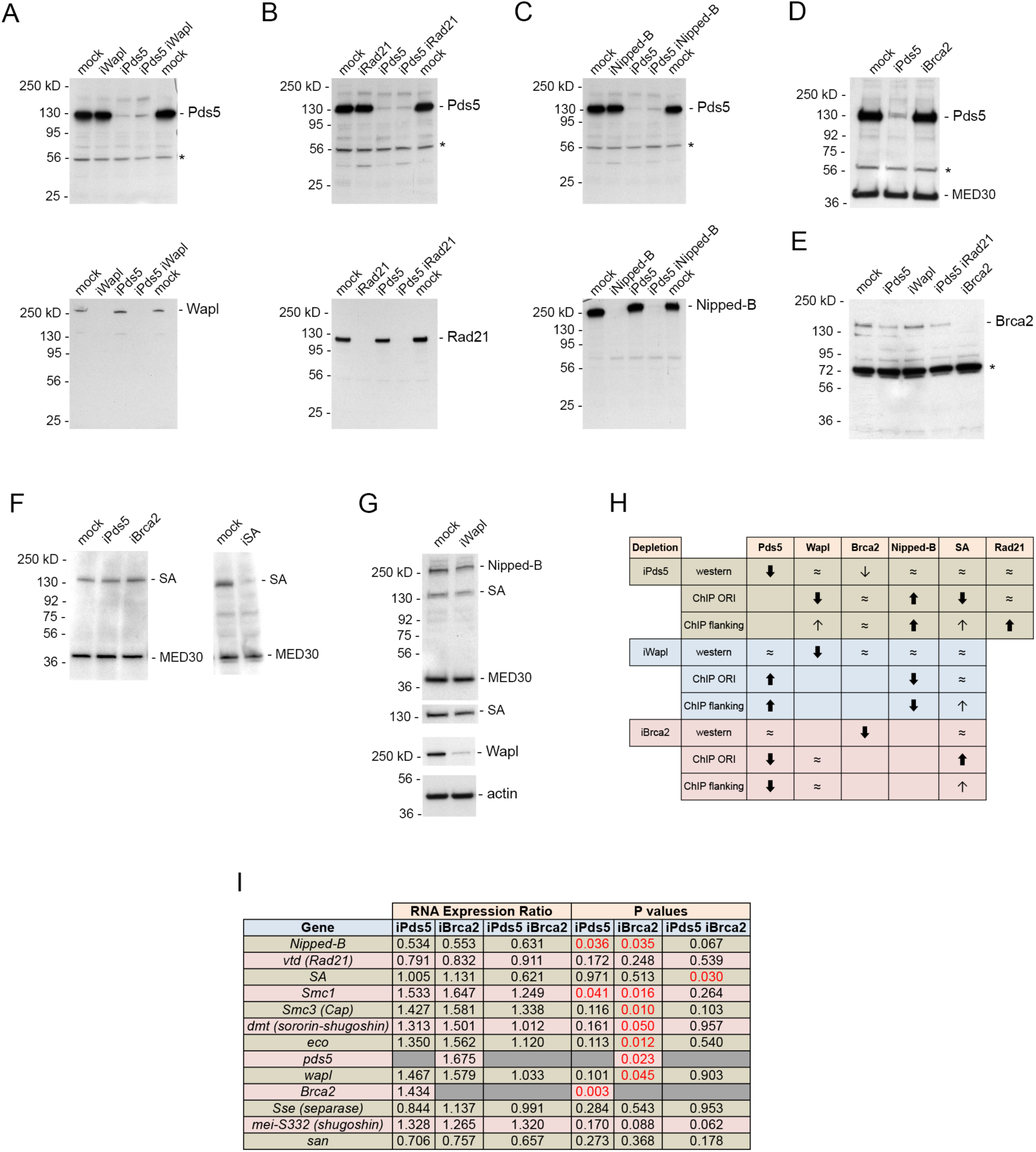
Effects of cohesion protein depletions on cohesion protein and transcripts. (**A**) The top and bottom panels are matched western blots of whole cell extracts of cells after the indicated single and double RNAi treatments for 4 to 5 days. The top panel is probed with anti-Pds5 antibody and the bottom panel is probed with anti-Wapl. The asterisk (*) indicates a non-specific band recognized by the Pds5 antibody used as a loading control. (**B**) Matched western blots with extracts from cells with the indicated single and double RNAi treatments. The top panel was probed with anti-Pds5 and the bottom panel was probed with anti-Rad21. (**C**) Matched western blots with extracts from cells with the indicated RNAi treatments. The top was probed with anti-Pds5 and the bottom was probed with anti-Nipped-B. (**D**) Western of extracts of cells after the indicated RNAi treatments (mock, iPds5, iBrca2) probed with anti-Pds5 and anti-MED30 as a loading control. (**E**) Western of extracts of cells after the indicated RNAi treatments (mock, iPds5, iWapl, iPds5 iRad21, iBrca2) treated probed with anti-Brca2. The asterisk (*) indicates a non-specific band recognized by anti-Brca2. (**F**) The left western shows extracts of cells with the indicated RNAi treatments (mock, iPds5, iBrca2) probed with anti-SA, and anti-MED30 Mediator subunit. The right panel is a western of extracts from mock-treated cells and cells depleted for SA (iSA) and anti-MED30 to demonstrate SA antibody specificity. (**G**) The top western shows extracts from mock-treated cells and cells depleted for Wapl (iWapl) probed with anti-Nipped-B, anti-SA, and anti-MED30. The second panel down shows a longer exposure for SA from the same blot. The third panel down shows the same blot when re-probed with anti-Wapl, and the bottom panel when re-probed with anti-actin. (**H**) Summary of the effects of Pds5, Wapl and Brca2 depletions (iPds5, iWapl, iBrca2) on the levels of the indicated proteins by western blot of whole cell extracts, and ChIP-seq enrichment at replication origin centers (ChIP ORI) or in regions flanking replications origins (ChIP flanking). “≈” indicates no significant change, thick down arrows indicate a large decrease, thin arrows indicate a small decrease, thick up arrows indicate a large increase, and thin up arrows indicate a small increase. Other than large decreases in the protein targeted by the RNAi treatment, the only noticeable effect of an RNAi treatment on a non-target protein is a small decrease in Brca2 with Pds5 depletion. See panel E for example westerns. There was no significant change in Brca2 ChIP-seq enrichment with Pds5 depletion. (**I**) Effects of Pds5, Brca2, and Pds5-Brca2 double depletion on cohesion factor transcripts measured by RNA-seq. The RNA Expression Ratio is the ratio of the level of the transcripts in depleted cells to the level in the mock-treated control cells. Gray boxes indicate where the double-stranded RNA used for RNAi treatment is detected by RNA-seq, preventing transcript quantification. Significant p values are in red. All expression comparisons shown gave q values greater than 0.05 (S8 Table).

**S2 Fig.**
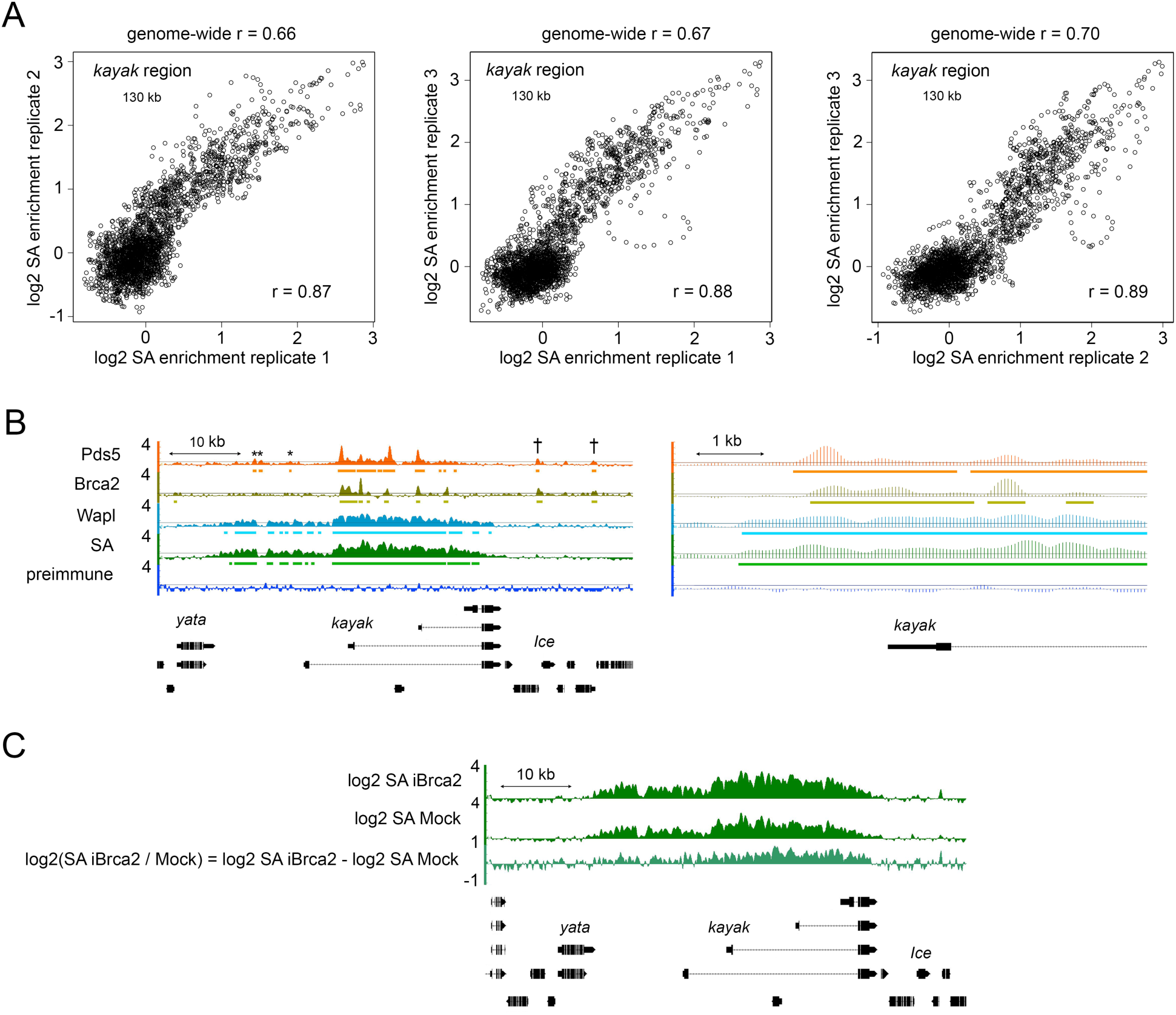
Examples of correlations between ChIP-seq biological replicates, preimmune ChIP-seq control, and calculating fold-changes in ChIP-seq enrichment. **(A)** SA ChIP-seq enrichment normalized to input chromatin (>45X genome coverage) every 50 bp across a 130 kilobase *kayak* region from three independent biological replicate experiments sequenced to at least 10X genome coverage are plotted against each other as examples of the reproducibility of the ChIP-seq method used for these studies. The genome-wide Pearson correlations between the two replicates plotted in each panel are above the plot, and the correlations in the 130 kilobase region surrounding *kayak* are given in the plot. (**B**) Genome browser views of Pds5, Brca2, Wapl, SA and preimmune serum ChIP-seq enrichment (log2 values) are shown as an example of the lack of significant enrichment with preimmune serum, indicating a lack of methodological artifacts. Bars underneath the ChIP-seq enrichment plots indicate where enrichment is in the 95 percentile or higher for at least 300 base pairs. Asterisks (*) indicate Pds5 binding sites without significant Brca2 occupancy. Daggers (†) indicate Pds5 – Brca2 binding sites in regions with little cohesin or Wapl. The right panel shows a higher resolution view of one of the active kayak gene promoters, illustrating the ChIP-seq enrichment values every 50 base pairs, simplifying downstream data analysis. (**C**) Example of an increase in SA enrichment at the kayak locus upon Brca2 depletion (iBrca2). The method used to calculate the fold-change in enrichment every 50 base pairs is the bottom track.

**S3 File. Locations of strongest early DNA replication origins in BG3 cells**

**S4 Fig.**
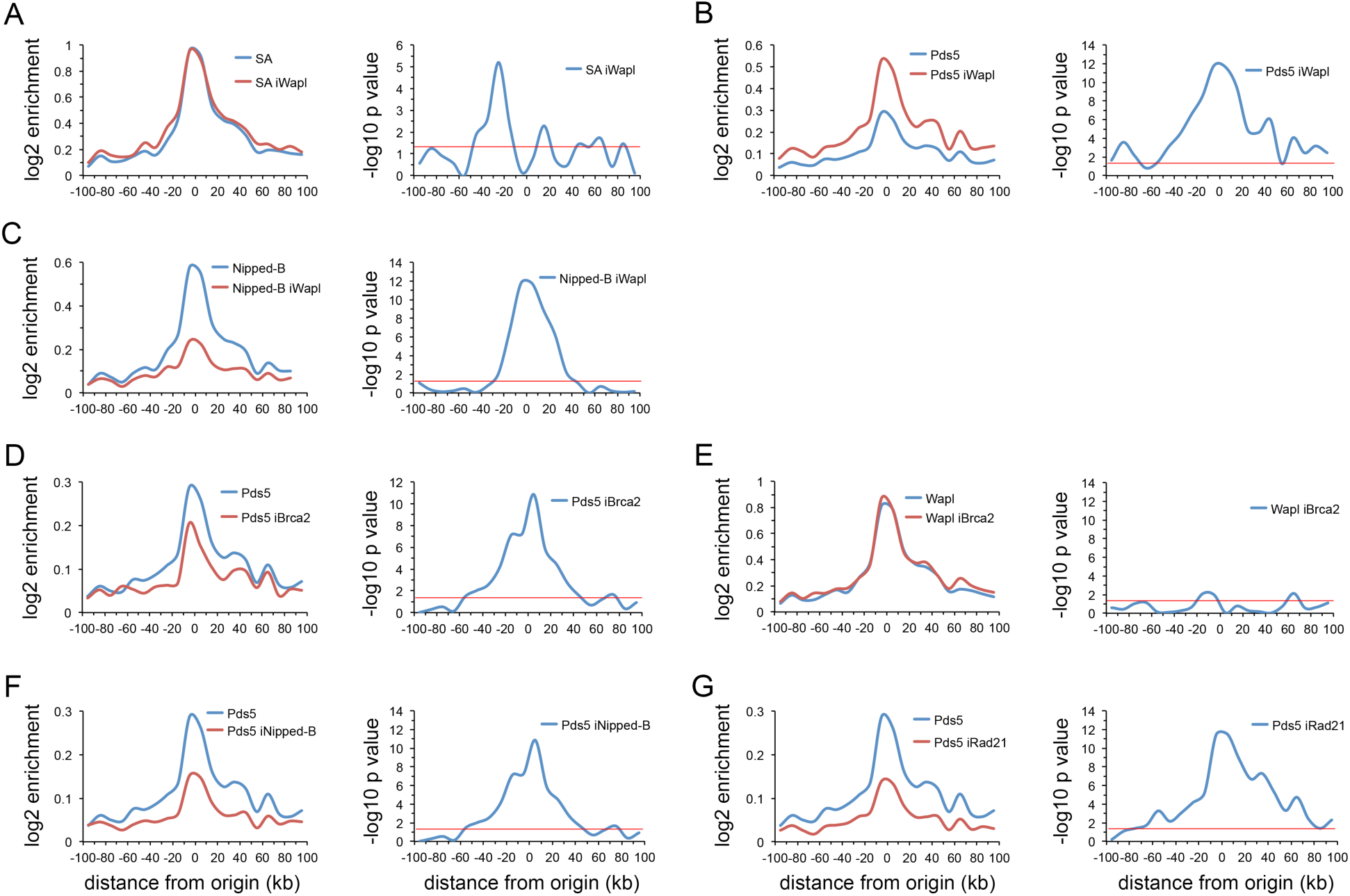
Meta-origin analyses in BG3 cells after Wapl, Brca2, Nipped-B and Rad21 depletion. (**A**) Left panel is the SA distribution in mock-treated control cells (blue, SA) and cells depleted for Wapl (red, SA iWapl). Right panel is the -log10 p values of each bin for the difference in control versus the depletion calculated using the Wilcoxon signed rank test. (**B**) Same as A for the Pds5 distribution. (**C**) Same as A for the Nipped-B distribution. (**D**) Left panel is the Pds5 meta-origin distribution in control (blue, Pds5) cells and cells depleted for Brca2 (red, Pds5 iBrca2). Right panel shows the –log10 p values. (**E**) Same as D for Wapl distribution. (**F**) Left panel is the Pds5 distribution in control (blue, Pds5) cells, and cells depleted for Nipped-B (red, Pds5 iNipped-B). Right panel shows the -log10 p values. (**G**) Same as F except for Rad21 depleted cells.

**S5 Fig.**
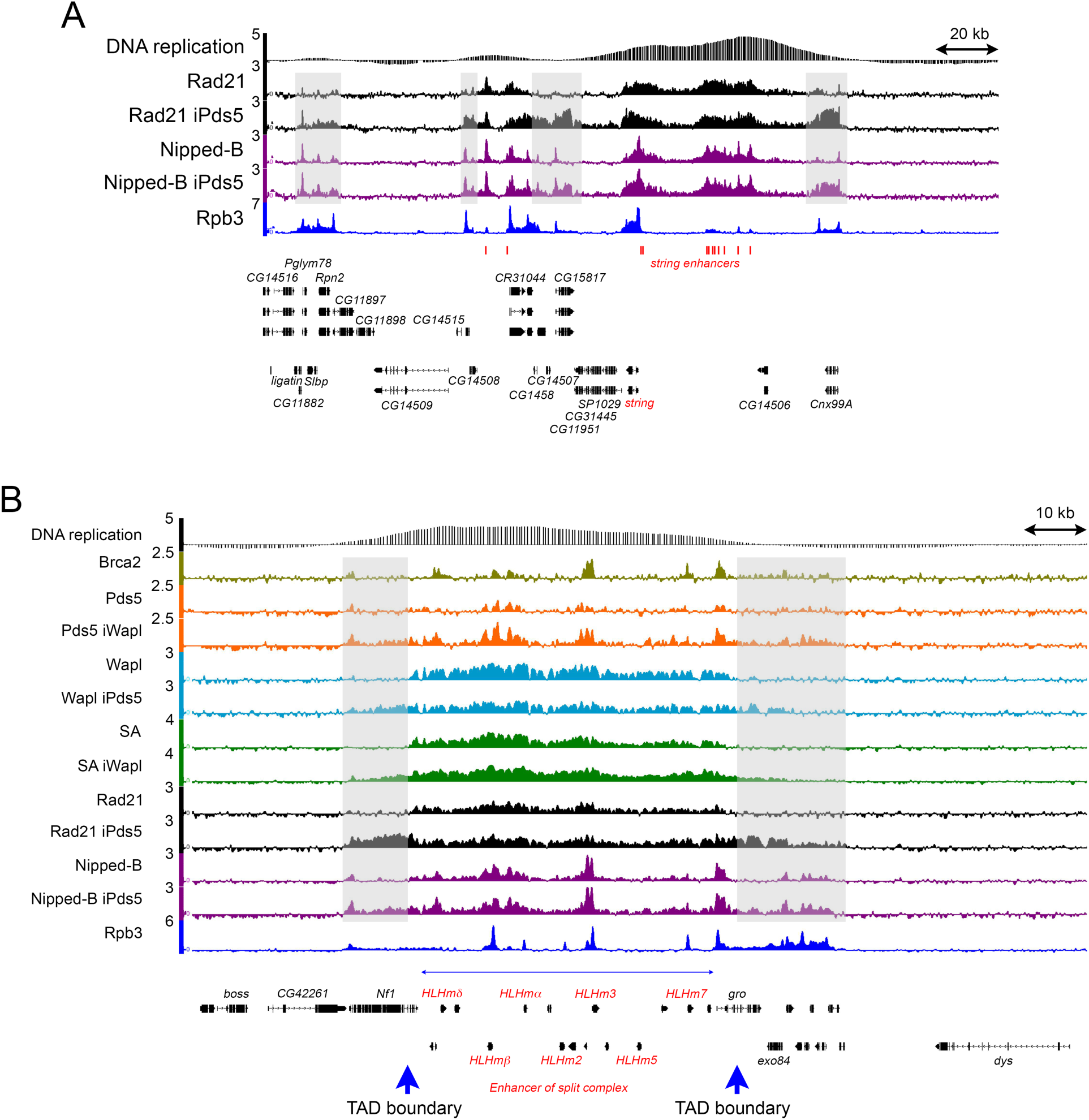
Examples of extended cohesin domains upon Pds5 or Wapl depletion. (**A**) Genome browser view of the *string* region showing skipping of spreading cohesin over inactive genes and intergenic regions. The *string* (*Cdc25*) gene and its active enhancers are labeled in red type. The top track is the early DNA replication pattern. The ChIP-seq tracks show the Rad21 (black) and Nipped-B (purple) binding patterns in control cells and cells depleted for Pds5 (iPds5). The bottom track shows the Rpb3 RNA polymerase pattern (blue). Shaded areas indicate regions of increased Rad21 and Nipped-B occupancy in Pds5-depleted cells. Pds5 depletion increases Rad21 and Nipped-B occupancy at the active gene cluster containing *Pglym78* some 100 kb centromere-proximal (left) to the origin, but not with inactive genes such as *CG14509* in the intervening region. Similarly, cohesin and Nipped-B increase substantially at the active *Cxn99A* gene to the right of the origin, but show only a modest increase in the intervening intergenic region. *CG15817* is active but binds much less cohesin than some active genes located further from the origin. Pds5 depletion increases cohesin and Nipped-B at *CG15817*. (**B**) Enhancer of split gene complex showing expansion of cohesin domains beyond TAD (topologically associating domain) boundaries upon Pds5 or Wapl depletion. Enhancer of split genes are labeled in red, and the thin blue horizontal arrow just above the genes shows the extent of the gene complex. Wide vertical blue arrows below the genes indicate the TAD boundaries determined by 3C (chromosome conformation capture) [36]. The top track shows the early DNA replication pattern. The ChIP-seq tracks show the Pds5 (orange) pattern in control cells and cells depleted for Wapl (iWapl), the Wapl pattern (light blue) in control cells and cells depleted for Pds5 (iPds5), the SA (green) binding in control cells and cells depleted for Wapl (iWapl), the Rad21 (black) binding in control cells and cells depleted for Pds5 (iPds5), the Nipped-B (purple) pattern in control cells and cells depleted for Pds5 (iPds5), and the Rpb3 (blue) RNA polymerase binding in control cells. Shaded areas indicate regions with increased occupancy by cohesion proteins upon Pds5 or Wapl depletion.

**S6 Fig.**
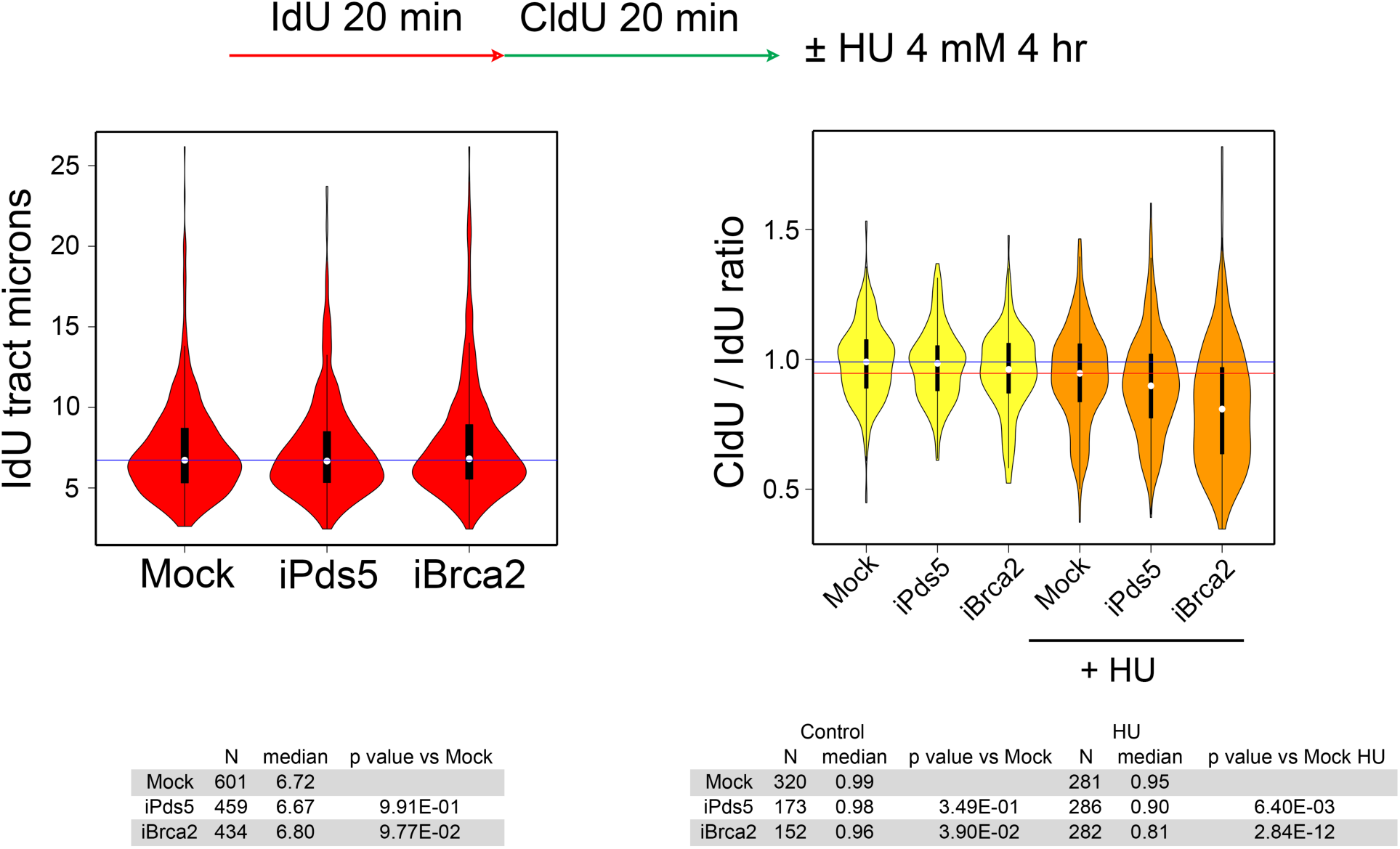
DNA fiber replication analysis in BG3 cells depleted for Pds5 and Brca2. The top diagram shows the labeling scheme and the hydroxyurea (HU) treatment after CldU incorporation. The left panel shows the lengths of the IdU tracts in Mock control cells, and cells depleted for Pds5 (iPds5) or Brca2 (iBrca2). The blue line indicates the median tract length in Mock control cells. The IdU tracts were combined for the HU-treated and untreated cells, and only IdU tracts continuous with CldU tracts were measured. The table beneath the left panel gives the median tract lengths in microns for each group and the p values using the Wilcoxon test. The right panel shows the ratio of lengths of the CldU tracts to the connected IdU tracts. The blue line indicates the median ratio in Mock control cells not treated with HU, and the red line indicates the median ratio in Mock control cells treated with HU. The table beneath the right panel gives the median ratio for all groups and the p values for the indicated comparisons using the Wilcoxon test. Similar results were obtained in two other independent experiments.

**S7 Table. Statistical tests of changes in cohesin and Nipped-B ChIP-seq enrichment at promoters, enhancers and PREs upon Pds5, Brca2 or Wapl depletion in BG3 cells**

**S8 Data. RNA-seq data in BG3 cells depleted for Nipped-B, Rad21, Pds5, Wapl, and Brca2, and control and brca2 mutant female 3^rd^ instar wing imaginal discs**

**S9 Fig.**
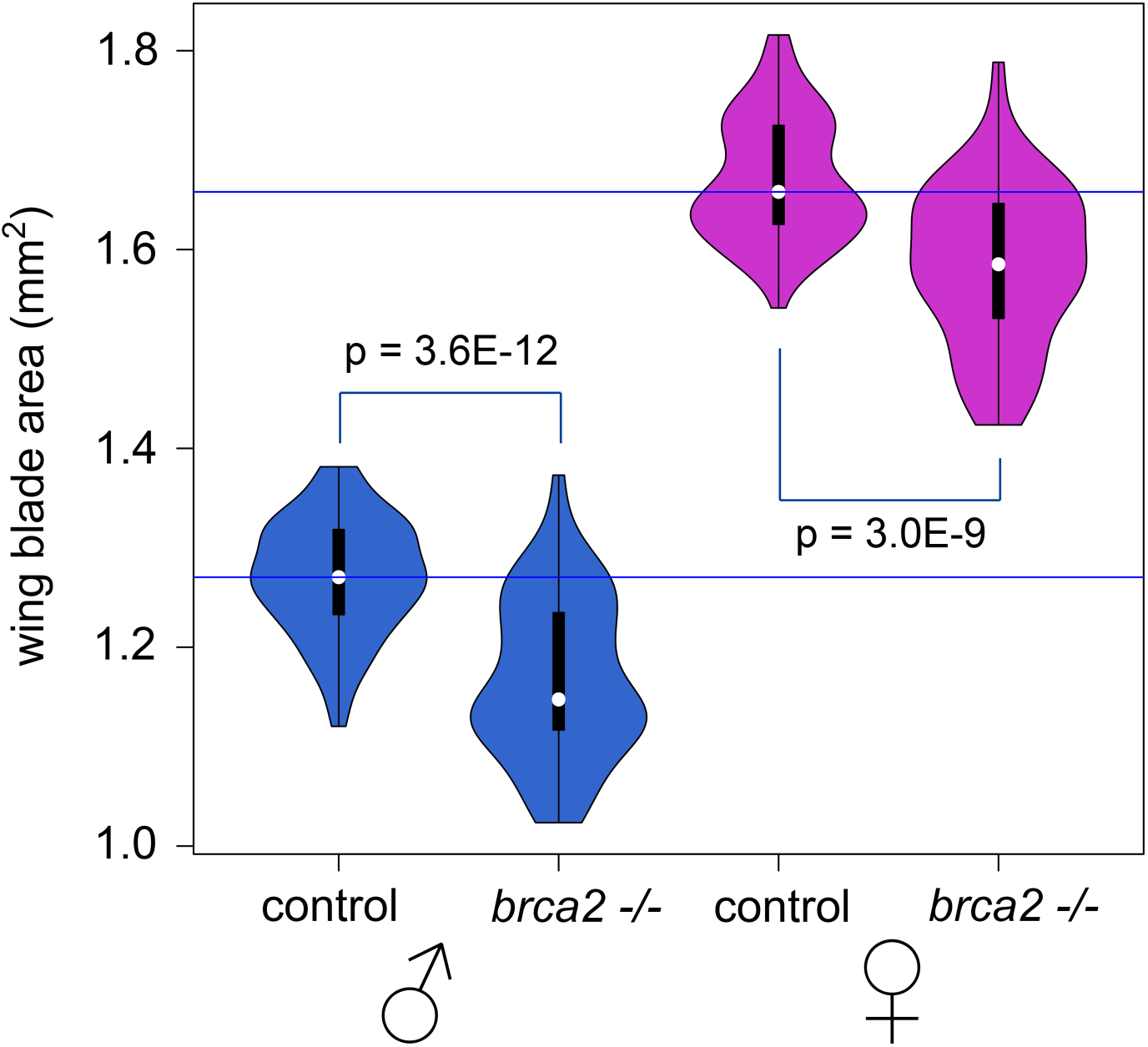
Adult wing areas in *brca2* mutant and control flies. Violin plots show the distributions of adult wing blade areas from the indicated groups of flies grown at 25°. The *brca2* −/− mutant wings are from ~20 homozygous *brca2*^*KO*^ and ~40 *brca2*^*KO*^ / *brca2*^*56e*^ flies and the control wings are from ~20 each from *cn bw*, Oregon R, *brca2*^*KO*^ / *cn bw* and *brca2*^*KO*^ / *cn bw* flies. The various mutant and control genotypes were combined to minimize the effects of genetic background differences that can influence growth. P values were calculated using the Wilcoxon test. Horizontal blue lines indicate the median wing areas for male and female control flies.

## References

1. Gligoris T, Löwe J. Structural insights into ring formation of cohesin and related Smc complexes. Trends Cell Biol. 2016;26: 680–93. doi: 10.1016/j.tcb.2016.04.002.

2. Ouyang Z, Yu H. Releasing the cohesin ring: A rigid scaffold model for opening the DNA exit gate by Pds5 and Wapl. Bioessays. 2017;39. doi: 10.1002/bies.201600207.

3. Rankin S, Dawson DS. Recent advances in cohesin biology. F1000Res. 2016;5. pii: F1000 Faculty Rev-1909. doi: 10.12688/f1000research.8881.1.

4. Uhlmann F. SMC complexes: from DNA to chromosomes. Nat Rev Mol Cell Biol. 2016;17: 399–412. doi: 10.1038/nrm.2016.30.

5. Hartman T, Stead K, Koshland D, Guacci V. Pds5p is an essential chromosomal protein required for both sister chromatid cohesion and condensation in *Saccharomyces cerevisiae*. J Cell Biol. 2000;151: 613–26.

6. Panizza S, Tanaka T, Hochwagen A, Eisenhaber F, Nasmyth K. Pds5 cooperates with cohesin in maintaining sister chromatid cohesion. Curr Biol. 2000;10: 1557–64.

7. Dorsett D, Eissenberg JC, Misulovin Z, Martens A, Redding B, McKim K. Effects of sister chromatid cohesion proteins on *cut* gene expression during wing development in Drosophila. Development. 2005;132: 4743–53. doi: 10.1242/dev.02064.

8. Verni F, Gandhi R, Goldberg ML, Gatti M. Genetic and molecular analysis of *wings apart-like* (*wapl*), a gene controlling heterochromatin organization in *Drosophila melanogaster*. Genetics. 2000;154: 1693–710.

9. Gause M, Misulovin Z, Bilyeu A, Dorsett D. Dosage-sensitive regulation of cohesin chromosome binding and dynamics by Nipped-B, Pds5, and Wapl. Mol Cell Biol. 2010;30: 4940–51. doi: 10.1128/MCB.00642-10.

10. Brough R, Bajrami I, Vatcheva R, Natrajan R, Reis-Filho JS, Lord CJ, Ashworth A. APRIN is a cell cycle specific BRCA2-interacting protein required for genome integrity and a predictor of outcome after chemotherapy in breast cancer. EMBO J. 2012;31: 1160–76. doi: 10.1038/emboj.2011.490.

11. Kusch T. Brca2-Pds5 complexes mobilize persistent meiotic recombination sites to the nuclear envelope. J Cell Sci. 2015;128: 717–27. doi: 10.1242/jcs.159988.

12. Couturier AM, Fleury H, Patenaude AM, Bentley VL, Rodrigue A, Coulombe Y, et al. Roles for APRIN (PDS5B) in homologous recombination and in ovarian cancer prediction. Nucleic Acids Res. 2016;44: 10879–97. doi: 10.1093/nar/gkw921.

13. Ouyang Z, Zheng G, Tomchick DR, Luo X, Yu H. Structural basis and IP6 requirement for Pds5-dependent cohesin dynamics. Mol Cell. 2016;62: 248–59. doi: 10.1016/j.molcel.2016.02.033.

14. Schaaf CA, Misulovin Z, Sahota G, Siddiqui AM, Schwartz YB, Kahn TG, et al. Regulation of the Drosophila Enhancer of split and *invected-engrailed* gene complexes by sister chromatid cohesion proteins. PLoS One 2009;4: e6202. doi: 10.1371/journal.pone.0006202.

15. Nishiyama T, Ladurner R, Schmitz J, Kreidl E, Schleiffer A, Bhaskara V, et al. Sororin mediates sister chromatid cohesion by antagonizing Wapl. Cell. 2010;143: 737–49. doi: 10.1016/j.cell.2010.10.031.

16. Yamada T, Tahara E, Kanke M, Kuwata K, Nishiyama T. Drosophila Dalmatian combines sororin and shugoshin roles in establishment and protection of cohesion. EMBO J. 2017;36: 1513–27. doi: 10.15252/embj.201695607.

17. Swain A, Misulovin Z, Pherson M, Gause M, Mihindukulasuriya K, Rickels RA, et al. Drosophila TDP-43 RNA-Binding protein facilitates association of sister chromatid cohesion proteins with genes, enhancers and Polycomb Response Elements. PLoS Genet. 2016;12:e1006331. doi: 10.1371/journal.pgen.1006331.

18. Dorsett D, Misulovin Z. Measuring sister chromatid cohesion protein genome occupancy in *Drosophila melanogaster* by ChIP-seq. Methods Mol Biol. 2017;1515: 125–139. doi: 10.1007/978-1-4939-6545-8_8.

19. Rowland BD, Roig MB, Nishino T, Kurze A, Uluocak P, Mishra A, et al. Building sister chromatid cohesion: smc3 acetylation counteracts an antiestablishment activity. Mol Cell. 2009;33: 763–74. doi: 10.1016/j.molcel.2009.02.028.

20. Shintomi K, Hirano T. Releasing cohesin from chromosome arms in early mitosis: opposing actions of Wapl-Pds5 and Sgo1. Genes Dev. 2009;23: 2224–36. doi: 10.1101/gad.1844309.

21. Sutani T, Kawaguchi T, Kanno R, Itoh T, Shirahige K. Budding yeast Wpl1(Rad61)-Pds5 complex counteracts sister chromatid cohesion-establishing reaction. Curr Biol. 2009;19: 492–7. doi: 10.1016/j.cub.2009.01.062.

22. Ouyang Z, Zheng G, Song J, Borek DM, Otwinowski Z, Brautigam CA, et al. Structure of the human cohesin inhibitor Wapl. Proc Natl Acad Sci U S A. 2013;110: 11355–60. doi: 10.1073/pnas.1304594110.

23. Eichinger CS, Kurze A, Oliveira RA, Nasmyth K. Disengaging the Smc3/kleisin interface releases cohesin `from Drosophila chromosomes during interphase and mitosis. EMBO J. 2013;32: 656–65. doi: 10.1038/emboj.2012.346.

24. Murayama Y, Uhlmann F. DNA entry into and exit out of the cohesin ring by an interlocking gate mechanism. Cell. 2015;163: 1628–40. doi: 10.1016/j.cell.2015.11.030.

25. Beckouët F, Srinivasan M, Roig MB, Chan KL, Scheinost JC, Batty P, et al. Releasing activity disengages cohesin's Smc3/Scc1 Interface in a process blocked by acetylation. Mol Cell. 2016;61: 563–574. doi: 10.1016/j.molcel.2016.01.026.

26. MacAlpine HK, Gordân R, Powell SK, Hartemink AJ, MacAlpine DM. Drosophila ORC localizes to open chromatin and marks sites of cohesin complex loading. Genome Res. 2010;20: 201–11. doi: 10.1101/gr.097873.109.

27. Gillespie PJ, Hirano T. Scc2 couples replication licensing to sister chromatid cohesion in Xenopus egg extracts. Curr Biol. 2004;14: 1598–603. doi: 10.1016/j.cub.2004.07.053.

28. Takahashi TS, Yiu P, Chou MF, Gygi S, Walter JC. Recruitment of Xenopus Scc2 and cohesin to chromatin requires the pre-replication complex. Nat Cell Biol. 2004;6: 991–6. doi: 10.1038/ncb1177.

29. Bermudez VP, Farina A, Higashi TL, Du F, Tappin I, Takahashi TS, Hurwitz J. In vitro loading of human cohesin on DNA by the human Scc2-Scc4 loader complex. Proc Natl Acad Sci U S A. 2012;109: 9366–71. doi: 10.1073/pnas.1206840109.

30. Takahashi TS, Basu A, Bermudez V, Hurwitz J, Walter JC. Cdc7-Drf1 kinase links chromosome cohesion to the initiation of DNA replication in Xenopus egg extracts. Genes Dev. 2008;22: 1894–905. doi: 10.1101/gad.1683308.

31. Guillou E, Ibarra A, Coulon V, Casado-Vela J, Rico D, Casal I, et al. Cohesin organizes chromatin loops at DNA replication factories. Genes Dev. 2010;24: 2812–22. doi: 10.1101/gad.608210.

32. Tittel-Elmer M, Lengronne A, Davidson MB, Bacal J, François P, Hohl M, et al. Cohesin association to replication sites depends on rad50 and promotes fork restart. Mol Cell. 2012;48: 98–108. doi: 10.1016/j.molcel.2012.07.004.

33. Pherson M, Misulovin Z, Gause M, Mihindukulasuriya K, Swain A, Dorsett D. Polycomb Repressive Complex 1 modifies transcription of active genes. Sci Adv. 2017;3: e1700944. doi: 10.1126/sciadv.1700944.

34. Misulovin Z, Schwartz YB, Li XY, Kahn TG, Gause M, MacArthur S, et al. Association of cohesin and Nipped-B with transcriptionally active regions of the *Drosophila melanogaster* genome. Chromosoma. 2008;117: 89–102. doi: 10.1007/s00412-007-0129-1.

35. Schaaf CA, Kwak H, Koenig A, Misulovin Z, Gohara DW, Watson A, et al. Genome-wide control of RNA polymerase II activity by cohesin. PLoS Genet. 2013;9: e1003382. doi: 10.1371/journal.pgen.1003382.

36. Schaaf CA, Misulovin Z, Gause M, Koenig A, Dorsett D. The Drosophila enhancer of split gene complex: architecture and coordinate regulation by notch, cohesin, and polycomb group proteins. G3 (Bethesda). 2013;3: 1785–94. doi: 10.1534/g3.113.007534.

37. Haarhuis JHI, van der Weide RH, Blomen VA, Yáñez-Cuna JO, Amendola M, van Ruiten MS, et al. The cohesin release factor WAPL restricts chromatin loop extension. Cell. 2017;169: 693–707.e14. doi: 10.1016/j.cell.2017.04.013.

38. Eagen KP, Aiden EL, Kornberg RD. Polycomb-mediated chromatin loops revealed by a subkilobase-resolution chromatin interaction map. Proc Natl Acad Sci U S A. 2017;114:8764–69. doi: 10.1073/pnas.1701291114.

39. Mourad R, Cuvier O. Computational identification of genomic features that influence 3D chromatin domain formation. PLoS Comput Biol. 2016;12: e1004908. doi: 10.1371/journal.pcbi.1004908.

40. Ulianov SV, Khrameeva EE, Gavrilov AA, Flyamer IM, Kos P, Mikhaleva EA, et al. Active chromatin and transcription play a key role in chromosome partitioning into topologically associating domains. Genome Res. 2016;26: 70–84. doi: 10.1101/gr.196006.

41. Kanke M, Tahara E, Huis In’t Veld PJ, Nishiyama T. Cohesin acetylation and Wapl-Pds5 oppositely regulate translocation of cohesin along DNA. EMBO J. 2016;35: 2686–2698. doi: 10.15252/embj.201695756.

42. Schlacher K, Christ N, Siaud N, Egashira A, Wu H, Jasin M. Double-strand break repair-independent role for BRCA2 in blocking stalled replication fork degradation by MRE11. Cell. 2011;145: 529–42. doi: 10.1016/j.cell.2011.03.041.

43. Kulemzina I, Schumacher MR, Verma V, Reiter J, Metzler J, Failla AV, et al. Cohesin rings devoid of Scc3 and Pds5 maintain their stable association with the DNA. PLoS Genet. 2012;8: e1002856. doi: 10.1371/journal.pgen.

44. Zhang N, Kuznetsov SG, Sharan SK, Li K, Rao PH, Pati D. A handcuff model for the cohesin complex. J Cell Biol. 2008;183: 1019–31. doi: 10.1083/jcb.200801157.

45. Zhang N, Pati D. Handcuff for sisters: a new model for sister chromatid cohesion. Cell Cycle. 2009;8: 399–402. doi: 10.4161/cc.8.3.7586.

46. Neuwald AF, Hirano T. HEAT repeats associated with condensins, cohesins, and other complexes involved in chromosome-related functions. Genome Res. 2000;10: 1445–52.

47. Kikuchi S, Borek DM, Otwinowski Z, Tomchick DR, Yu H. Crystal structure of the cohesin loader Scc2 and insight into cohesinopathy. Proc Natl Acad Sci U S A. 2016;113: 12444–9. doi: 10.1073/pnas.1611333113.

48. Chao WC, Murayama Y, Muñoz S, Jones AW, Wade BO, Purkiss AG, et al. Structure of the cohesin loader Scc2. Nat Commun. 2017;8: 13952. doi: 10.1038/ncomms13952.

49. Lee BG, Roig MB, Jansma M, Petela N, Metson J, Nasmyth K, Löwe J. Crystal structure of the cohesin gatekeeper Pds5 and in complex with kleisin Scc1. Cell Rep. 2016;14: 2108–15. doi: 10.1016/j.celrep.2016.02.020.

50. Wells JN, Gligoris TG, Nasmyth KA, Marsh JA. Evolution of condensin and cohesin complexes driven by replacement of Kite by Hawk proteins. Curr Biol. 2017;27: R17–R18. doi: 10.1016/j.cub.2016.11.050.

51. Carretero M, Ruiz-Torres M, Rodríguez-Corsino M, Barthelemy I, Losada A. Pds5B is required for cohesion establishment and Aurora B accumulation at centromeres. EMBO J. 2013;32: 2938–49. doi: 10.1038/emboj.2013.230.

52. Muir KW, Kschonsak M, Li Y, Metz J, Haering CH, Panne D. Structure of the Pds5-Scc1 complex and implications for cohesin function. Cell Rep. 2016;14:2116–26. doi: 10.1016/j.celrep.2016.01.078.

53. Schaaf CA, Misulovin Z, Gause M, Koenig A, Gohara DW, Watson A, Dorsett D. 2013. Cohesin and polycomb proteins functionally interact to control transcription at silenced and active genes. PLoS Genet. 2013;9: e1003560. doi: 10.1371/journal.pgen.1003560.

54. Fay A, Misulovin Z, Li J, Schaaf CA, Gause M, Gilmour DS, Dorsett D. Cohesin selectively binds and regulates genes with paused RNA polymerase. Curr Biol. 2011;21: 1624–34. doi: 10.1016/j.cub.2011.08.036.

55. Klovstad M, Abdu U, Schüpbach T. Drosophila brca2 is required for mitotic and meiotic DNA repair and efficient activation of the meiotic recombination checkpoint. PLoS Genet. 2008;4: e31. doi: 10.1371/journal.pgen.0040031.

56. Wu Y, Gause M, Xu D, Misulovin Z, Schaaf CA, Mosarla RC, et al. Drosophila *Nipped-B* mutants model Cornelia de Lange syndrome in growth and behavior. PLoS Genet. 2015;11: e1005655. doi: 10.1371/journal.pgen.1005655.

57. Busslinger GA, Stocsits RR, van der Lelij P, Axelsson E, Tedeschi A, Galjart N, Peters JM. Cohesin is positioned in mammalian genomes by transcription, CTCF and Wapl. Nature. 2017;544: 503–7. doi: 10.1038/nature22063.

58. Xiao T, Wallace J, Felsenfeld G. Specific sites in the C terminus of CTCF interact with the SA2 subunit of the cohesin complex and are required for cohesin-dependent insulation activity. Mol Cell Biol. 2011;31: 2174–83. doi: 10.1128/MCB.05093-11.

59. Glynn EF, Megee PC, Yu HG, Mistrot C, Unal E, Koshland DE, et al. Genome-wide mapping of the cohesin complex in the yeast *Saccharomyces cerevisiae*. PLoS Biol. 2004;2: E259.

60. Lengronne A, Katou Y, Mori S, Yokobayashi S, Kelly GP, Itoh T, et al. Cohesin relocation from sites of chromosomal loading to places of convergent transcription. Nature. 2004;430: 573–8.

61. Ocampo-Hafalla M, Muñoz S, Samora CP, Uhlmann F. Evidence for cohesin sliding along budding yeast chromosomes. Open Biol. 2016;6. pii: 150178. doi: 10.1098/rsob.150178.

62. Skibbens RV. Establishment of sister chromatid cohesion. Curr Biol. 2009;19: R1126–32. doi: 10.1016/j.cub.2009.10.067.

63. Tong K, Skibbens RV. Pds5 regulators segregate cohesion and condensation pathways in *Saccharomyces cerevisiae*. Proc Natl Acad Sci U S A. 2015;112: 7021–6. doi: 10.1073/pnas.1501369112.

64. Chatterjee A, Zakian S, Hu XW, Singleton MR. Structural insights into the regulation of cohesion establishment by Wpl1. EMBO J. 2013;32: 677–87. doi: 10.1038/emboj.2013.16.

65. Roig MB, Löwe J, Chan KL, Beckouët F, Metson J, Nasmyth K. Structure and function of cohesin's Scc3/SA regulatory subunit. FEBS Lett. 2014;588: 3692–702. doi: 10.1016/j.febslet.2014.08.015.

66. Geck P, Sonnenschein C, Soto AM. The D13S171 marker, misannotated to BRCA2, links the AS3 gene to various cancers. Am J Hum Genet. 2001;69: 461–3. doi: 10.1086/321968.

67. Rollins RA, Morcillo P, Dorsett D. Nipped-B, a Drosophila homologue of chromosomal adherins, participates in activation by remote enhancers in the *cut* and *Ultrabithorax* genes. Genetics. 1999;152: 577–93.

68. Barrington C, Finn R, Hadjur S. Cohesin biology meets the loop extrusion model. Chromosome Res. 2017;25: 51–60. doi: 10.1007/s10577-017-9550-3.

69. Kagey MH, Newman JJ, Bilodeau S, Zhan Y, Orlando DA, van Berkum NL, et al. Mediator and cohesin connect gene expression and chromatin architecture. Nature. 2010 23;467: 430–5. doi: 10.1038/nature09380.

70. Krantz ID, McCallum J, DeScipio C, Kaur M, Gillis LA, Yaeger D, et al. Cornelia de Lange syndrome is caused by mutations in *NIPBL*, the human homolog of *Drosophila melanogaster Nipped-B*. Nat Genet. 2004;36: 631–5. doi: 10.1038/ng1364.

71. Tonkin ET, Wang TJ, Lisgo S, Bamshad MJ, Strachan T. *NIPBL*, encoding a homolog of fungal Scc2-type sister chromatid cohesion proteins and fly Nipped-B, is mutated in Cornelia de Lange syndrome. Nat Genet. 2004;36: 636–41. doi: 10.1038/ng1363.

72. Deardorff MA, Kaur M, Yaeger D, Rampuria A, Korolev S, Pie J, et al. Mutations in cohesin complex members SMC3 and SMC1A cause a mild variant of Cornelia de Lange syndrome with predominant mental retardation. Am J Hum Genet. 2007;80: 485–94. doi: 10.1086/511888.

73. Musio A, Selicorni A, Focarelli ML, Gervasini C, Milani D, Russo S, et al. X-linked Cornelia de Lange syndrome owing to *SMC1L1* mutations. Nat Genet. 2006;38: 528–30. doi: 10.1038/ng1779.

74. Deardorff MA, Bando M, Nakato R, Watrin E, Itoh T, Minamino M, et al. HDAC8 mutations in Cornelia de Lange syndrome affect the cohesin acetylation cycle. Nature. 2012;489: 313–7. doi: 10.1038/nature11316.

75. Deardorff MA, Wilde JJ, Albrecht M, Dickinson E, Tennstedt S, Braunholz D, et al. RAD21 mutations cause a human cohesinopathy. Am J Hum Genet. 2012;90: 1014–27. doi: 10.1016/j.ajhg.2012.04.019.

76. Dorsett D, Krantz ID. On the molecular etiology of Cornelia de Lange syndrome. Ann N Y Acad Sci. 2009;1151: 22–37. doi: 10.1111/j.1749-6632.2008.03450.x.

77. Liu J, Krantz ID. Cornelia de Lange syndrome, cohesin, and beyond. Clin Genet. 2009;76:303–14. doi: 10.1111/j.1399-0004.2009.01271.x.

78. Zhang B, Jain S, Song H, Fu M, Heuckeroth RO, Erlich JM, et al. Mice lacking sister chromatid cohesion protein PDS5B exhibit developmental abnormalities reminiscent of Cornelia de Lange syndrome. Development. 2007;134: 3191–201. doi: 10.1242/dev.005884.

79. Zhang B, Chang J, Fu M, Huang J, Kashyap R, Salavaggione E, et al. Dosage effects of cohesin regulatory factor PDS5 on mammalian development: implications for cohesinopathies. PLoS One. 2009;4: e5232. doi: 10.1371/journal.pone.0005232.

80. Kawauchi S, Calof AL, Santos R, Lopez-Burks ME, Young CM, Hoang MP, et al. Multiple organ system defects and transcriptional dysregulation in the Nipbl(+/-) mouse, a model of Cornelia de Lange Syndrome. PLoS Genet. 2009;5: e1000650. doi: 10.1371/journal.pgen.1000650.

81. Cunningham MD, Gause M, Cheng Y, Noyes A, Dorsett D, Kennison JA, Kassis JA. Wapl antagonizes cohesin binding and promotes Polycomb-group silencing in Drosophila. Development. 2012;139: 4172–9. doi: 10.1242/dev.084566.

82. Kennison JA, Tamkun JW. Dosage-dependent modifiers of polycomb and antennapedia mutations in Drosophila. Proc Natl Acad Sci U S A. 1988;85: 8136–40.

83. Hallson G, Syrzycka M, Beck SA, Kennison JA, Dorsett D, Page SL, et al. The Drosophila cohesin subunit Rad21 is a trithorax group (trxG) protein. Proc Natl Acad Sci U S A. 2008;105: 12405–10. doi: 10.1073/pnas.0801698105.

84. Hakem R, de la Pompa JL, Mak TW. Developmental studies of *Brca1* and *Brca2* knock-out mice. J Mammary Gland Biol Neoplasia. 1998;3: 431–45.

85. Connor F, Bertwistle D, Mee PJ, Ross GM, Swift S, Grigorieva E, et al. Tumorigenesis and a DNA repair defect in mice with a truncating Brca2 mutation. Nat Genet. 1997;17: 423–30. doi: 10.1038/ng1297-423

86. Howlett NG, Taniguchi T, Olson S, Cox B, Waisfisz Q, De Die-Smulders C, et al. Biallelic inactivation of *BRCA2* in Fanconi anemia. Science. 2002;297: 606–9.

87. Wagner JE, Tolar J, Levran O, Scholl T, Deffenbaugh A, Satagopan J, et al. Germline mutations in *BRCA2*: shared genetic susceptibility to breast cancer, early onset leukemia, and Fanconi anemia. Blood. 2004;103: 3226–9.

88. R Core Team (2013). R: A language and environment for statistical computing. R Foundation for Statistical Computing, Vienna, Austria. ISBN 3-900051-07-0, URL http://www.Rproject.org/.

89. Freese NH, Norris DC, Loraine AE. Integrated genome browser: visual analytics platform for genomics. Bioinformatics. 2016;32: 2089–95. doi: 10.1093/bioinformatics/btw069.

90. Jackson DA, Pombo A. Replicon clusters are stable units of chromosome structure: evidence that nuclear organization contributes to the efficient activation and propagation of S phase in human cells. J Cell Biol. 1998;140: 1285–95.

